# Integrating Multi-Modal Cancer Data Using Deep Latent Variable Path Modelling

**DOI:** 10.1101/2024.06.13.598616

**Authors:** Alex Ing, Alvaro Andrades, Marco Raffaele Cosenza, Jan O. Korbel

## Abstract

Cancers are commonly characterised by a complex pathology encompassing genetic, microscopic and macroscopic features, which can be probed individually using imaging and omics technologies. Integrating this data to obtain a full understanding of pathology remains challenging. We introduce a new method called Deep Latent Variable Path Modelling (DLVPM), which combines the representational power of deep learning with the capacity of path modelling to identify relationships between interacting elements in a complex system. To evaluate the capabilities of DLVPM, we initially trained a foundational model to map dependencies between SNV, Methylation, miRNA-Seq, RNA-Seq and Histological data using Breast Cancer data from The Cancer Genome Atlas (TCGA). This method exhibited superior performance in mapping associations between data types compared to classical path modelling. We additionally performed successful applications of the model to: stratify single-cell data, identify synthetic lethal interactions using CRISPR-Cas9 screens derived from cell-lines, and detect histologic-transcriptional associations using spatial transcriptomic data. Results from each of these data types can then be understood with reference to the same holistic model of illness.

## Introduction

Many common illnesses such as cancer, cardiovascular diseases, and neurological disorders result from complex pathologies possessing genetic, microscopic and macroscopic components^1–4^. Over recent decades, the invention and widespread use of diverse omics and imaging technologies has provided significant insights into the mechanisms underpinning these diseases^5^. However, analysed individually, these technologies may illuminate only a single aspect of the pathology. A comprehensive understanding of complex disease necessitates the integration of these disparate data types^6^. Current methods lack the ability to model and jointly incorporate both structured and unstructured data, as well as modelling direct and indirect relationships between these data types.

Cancers are characterised by a complex pathology. Cellular functions are dictated via multiple layers of genetic information and processing. In cancer, this information is corrupted, and normal processes are subverted in highly complex ways that give cancer cells the ability to survive, proliferate and metastasize^1^. Recent studies have revealed a diversity of somatic mutation classes, widespread epigenetic changes, and significant alterations in gene expression, all of which exhibit high heterogeneity across tumours, even within the same tissue^7^. Despite its molecular genesis, cancer is still primarily diagnosed and understood clinically through histological imaging; this process involves extracting, sectioning, staining, and imaging tumour biopsies to identify aberrations in tissue microstructure linked to specific clinical outcomes^8^. Efficient, integrative approaches that systematically combine multi-omic and imaging modalities could lead to deeper insights into cancer biology.

Deep learning methods excel at processing unstructured data, such as imaging, by identifying complex patterns without explicit programming^9^. In recent years, numerous hypothesis-driven studies have been conducted with a focus on predicting the presence of genetic characteristics of cancer, such as driver mutations and clinical status, using histological data^10,11^. Deep learning methods have also proved highly effective in modelling the complex interdependencies between genes in multi-omic datasets^12,13^. Other research efforts have aimed to integrate histological and genetic data to predict clinically significant outcomes, including patient survival times^14,15^. Despite these advancements, exploratory research that seeks to map the complex interactions between various layers of genetic information and histological data is still in its infancy, presenting a largely uncharted frontier in oncology. All-encompassing methods are required to comprehensively map the causal and statistical dependencies between different data-types relevant in cancer biology.

Path modelling, or structural equation modelling, is a powerful and widely used class of techniques utilized primarily in epidemiology^16^, social sciences^17^, and econometrics^18^. These methods can offer insights into relationships in complex systems and in testing theoretical and, in some cases, causal models^19,20^ by quantifying direct and indirect relationships between data-types. Nevertheless, these methods currently have limited representational power. Consequently, they share similar constraints with classical techniques used for classification and regression, which are inadequate for the evaluation of unstructured data, such as images, and the handling of complex, non-linear patterns frequently encountered in biological data^12,13^.

In this study, we introduce a Deep Learning based method for Path Modelling called Deep Latent Variable Path Modelling (DLVPM). This method combines the representational power of Deep Learning, with the ability of Path Modelling to map complex dependencies between data types. In the cancer context, this allows us to model genetic and epigenetic interactions with gene expression, which in turn result in the microscopically visible aberrations in tissue structure that are characteristic of cancer. A crucial strength of the method is its modular nature, which allows sub-models trained for each individual modality to be characterised further on new datasets.

We initially trained DLVPM on the TCGA (The Cancer Genome Atlas) breast cancer dataset^21^, one of the most comprehensive and well-annotated datasets combining imaging and multi-omics data modalities. Training on this dataset provides a robust foundational model, which we evaluated comprehensively on single cell, cell-line and spatial transcriptomics datasets. Insights gained from these different data-types can all be understood with reference to the same underlying foundational model.

This approach offers a holistic view of cancer, illustrating the power of DLVPM as a singular, comprehensive model for multi-layered data integration.

## Results

### Deep Latent Variable Path Modelling (DLVPM)

Path modelling/structural equation modelling methods are a family of procedures used for mapping dependencies between different data-types^19^. These methods are able to model arbitrarily many data-types simultaneously, providing a holistic view of a system of interacting elements. Path modelling analyses begin with the user specifying the path model itself. This model encodes hypotheses about the relationships between data types included in an analysis. These models are usually represented visually as a network graph (see Figure 1a), and mathematically as an adjacency matrix:

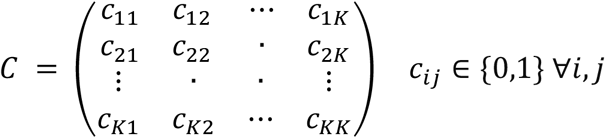

**Figure 1a:**
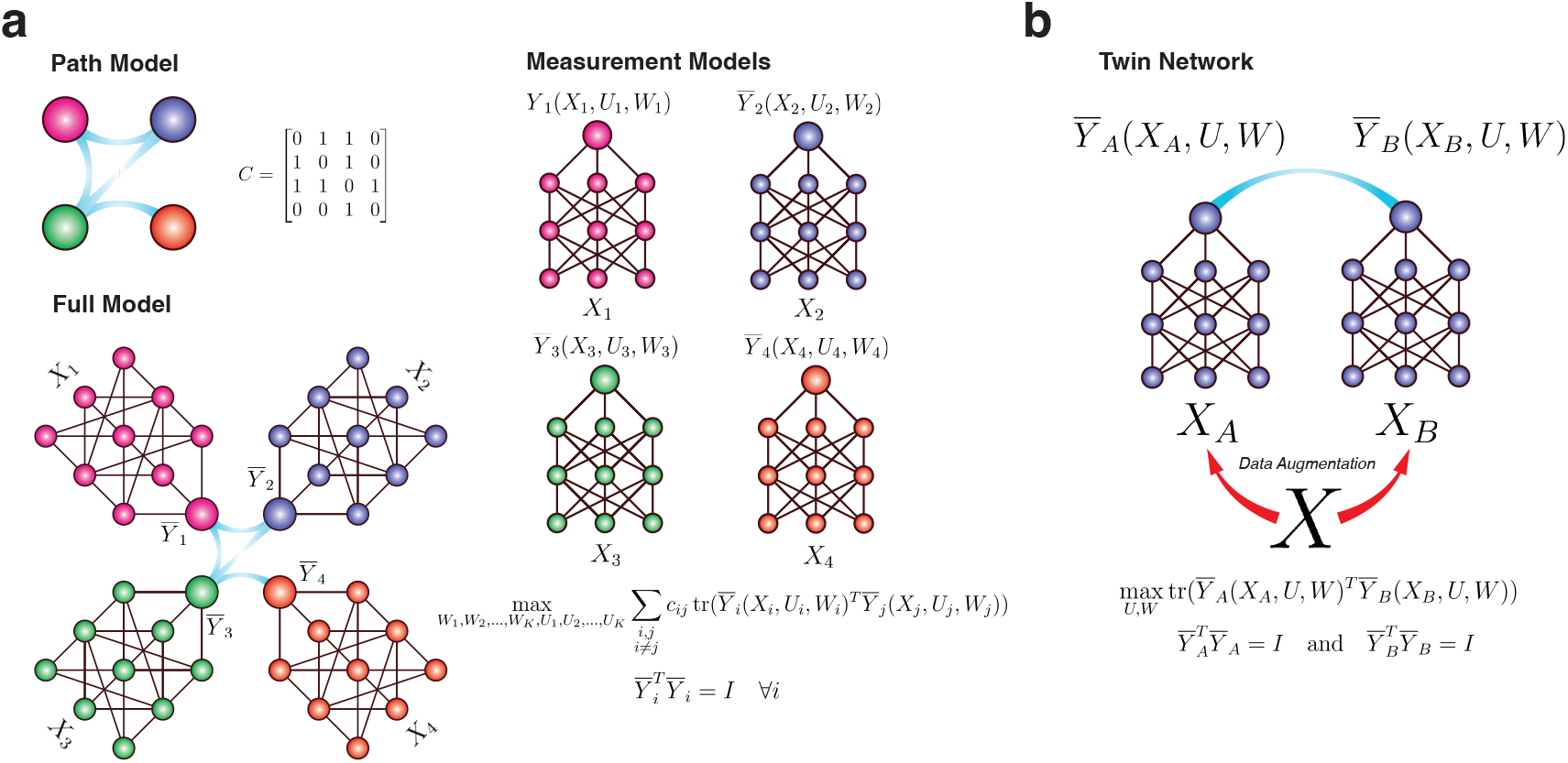
This figure shows the constituent parts of a DLVPM model. The path model defines the data-types that are connected to one another. Measurement models for each data-type are used to construct deep latent variables (DLVs) that are optimized to be strongly correlated between data-types. The overall model combines both the path model and the measurement models. This image represents DLVPM in a situation where four data types are available. b: Shows how DLVPM can be used in a Twin/Siamese network configuration. Here, augmented versions of the same input are fed to a network, and the network is trained to learn DLVs that are invariant to these augmentations.

The adjacency matrix is a square matrix where the elements *c*_*ij*_ represent connections between data types *i* and *j*, and *K* is the total number of data-types under analysis. Each element in the matrix indicates the presence (value of one) or absence (zero value) of a direct influence from one data-type to another.

In classical path modelling, techniques like Partial Least Squares Path Modelling (PLS-PM) are employed to derive Latent Variables (LVs) that exhibit optimal correlation among datasets linked by the path model. However, such techniques are limited to modelling linear effects^19^.

Deep neural networks excel in their ability to model non-linear effects, and to process structured and unstructured data. Most neural networks can be written in the general form:

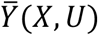

Where 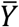 is the network output, *X* is some data-input and *U* is the set of network parameters, including weights, biases and other network parameters.

In DLVPM, we define a collection of sub-models, one for each data-type, indexed here by the subscript *i*:

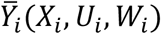

Where 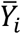 is the network output, a set of Deep Latent Variables (DLVs); *U*_*i*_ is the set of parameters up to the penultimate network layer, and the *W*_*i*_ term here corresponds to the network weights on the last layer of the neural network. This weight is displayed separately as it represents a linear projection and is critical to the way DLVPM is trained. These sub-models are called measurement models^19^.

The DLVPM algorithm is then trained to construct DLVs from each measurement model, which are optimized to be maximally associated with DLVs from other measurement models, connected by the path model. This optimization criteria can be written as:

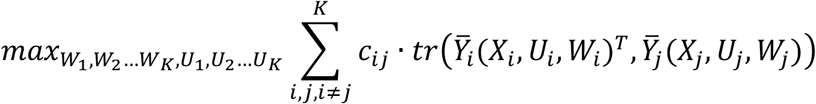

Where *c*_*ij*_ represents the association matrix input from data type *i* to data type *j*, and *tr* denotes the matrix trace. DLVs derived from each data-type are constrained to be orthogonal to one another:

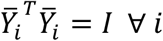

Where *I* is the identity matrix. These DLVs are then optimized to be strongly correlated across data types connected by the path model, while maintaining orthogonality within each data type, thus capturing the essence of each data type’s contribution to the system, whilst minimizing information redundancy within the model. Following model training, the DLVPM algorithm results in a set of orthogonal path models representing associations between DLVs constructed from each data type. In deep learning parlance, these DLVs can be considered to represent a joint embedding. DLVPM’s training process is both iterative and end-to-end, enabling the model to learn directly from raw data to the final output without the requirement for manual feature engineering.

The DLVPM method is extremely general. The measurement model formula, 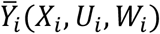, hides a high level of generality and complexity. In practice, almost any kind of neural network can be used here. This means that the method can be used to create embeddings shared by feedforward networks, convolutional networks, transformers etc., where architectural choices will depend on the data under analysis.

We introduce two different formulations of DLVPM, using different orthogonalization procedures. During training, the orthogonalization constraint is achieved via ZCA-whitening or iterative orthogonalization. ZCA-whitening is a widely used approach for whitening in deep learning. Iterative orthogonalization has the advantage that it prioritizes DLVs by their importance - a feature of considerable significance in the biological application presented here.

### DLVPM-Twins

Although DLVPM is primarily designed to uncover associations between multiple data types, it also excels at discovering useful representations of a single data type. In this context, DLVPM mirrors the objectives of confirmatory factor analysis (CFA)^22^ within classical path modelling—each serves to distil complex data into simpler, interpretable structures. However, while CFA confines itself to linear relationships, DLVPM extends this capacity into the non-linear domain by enabling the use of deep neural network architectures. When used in this manner, DLVPM falls into the class of methods called Siamese or Twin-networks^23^. This class of methods has become popular across a wide range of fields in recent years ^24,25^. Using this type of method, two augmented (distorted) versions of the same input are passed to a network. The model is then trained to learn features invariant to applied augmentations, thus promoting the development of robust and generalizable features (see Figure 1b). The optimization criteria for this method can be written as:

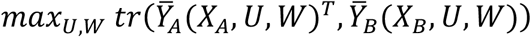

Subject to the constraint:

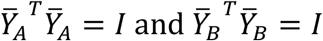

Where 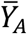 and 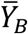 are outputs of the neural network with weights *U* and *W*. Here, *X*_*A*_ and *X*_*B*_ are different augmentations of the same input *X*. As was the case for the full DLVPM path modelling procedure, both ZCA-whitening and iterative orthogonalization schemes were used to impose the orthogonality.

A full and robust mathematical formulation of DLVPM is given in the Methods section. The algorithm is illustrated further in Extended Data Figure 1.

### Confounding Effects

Previous research has highlighted how factors like tumour purity and the acquisition site can undermine the replicability and generalizability of studies on molecular and histological data^26,27^. To address these issues, we developed a new approach for removing the effect of confounders; this method is implemented as a custom neural network layer. This layer uses the Moore-Penrose pseudo-inverse of a matrix of nuisance covariates to remove the effect of confounding variables. We used this method to remove confounding effects of site and tumour purity in all DLVPM analyses. Notably, this versatile layer represents a separate contribution from the main DLVPM method, and can be used in any neural network model. The mechanics of this approach are thoroughly detailed in the Methods and are illustrated in Extended Data Figure 2.

### Benchmarking DLVPM-Twins against other twin networks

DLVPM is, to our knowledge, a first in class method for path-modelling using neural networks. To establish its efficacy in learning meaningful data representations, we benchmarked DLVPM-Twins against other siamese/twin networks in the task of learning useful representations of a single data type – by performing testing of DLVPM-Twins against the VicReg^25^ and BarlowTwins^28^ methods.

Histological whole-slide images (WSIs) are generally in the gigapixel size range. Due to their vast size, these images are typically broken up into sub-images called tiles for further analysis^10^. This creates an ideal test case for twin networks, which excel in cases where we have more test examples than data labels: WSIs are made up of thousands of tiles. This is a great benefit in the context of deep learning, where having so many examples helps to prevent model overfitting.

Twin models are trained by passing the network augmented versions of the same data. In the present context, this means flipping, rotating, shearing and altering the colour of image tiles that are passed to the network. The model is then trained to learn orthogonal DLVs that are invariant to these augmentations. As meaningful outputs of the network should be invariant to these distortions, the network is encouraged to learn useful representations of the data (see Figure 2a). In this analysis, we used tiles extracted at x20 magnification.

**Figure 2a:**
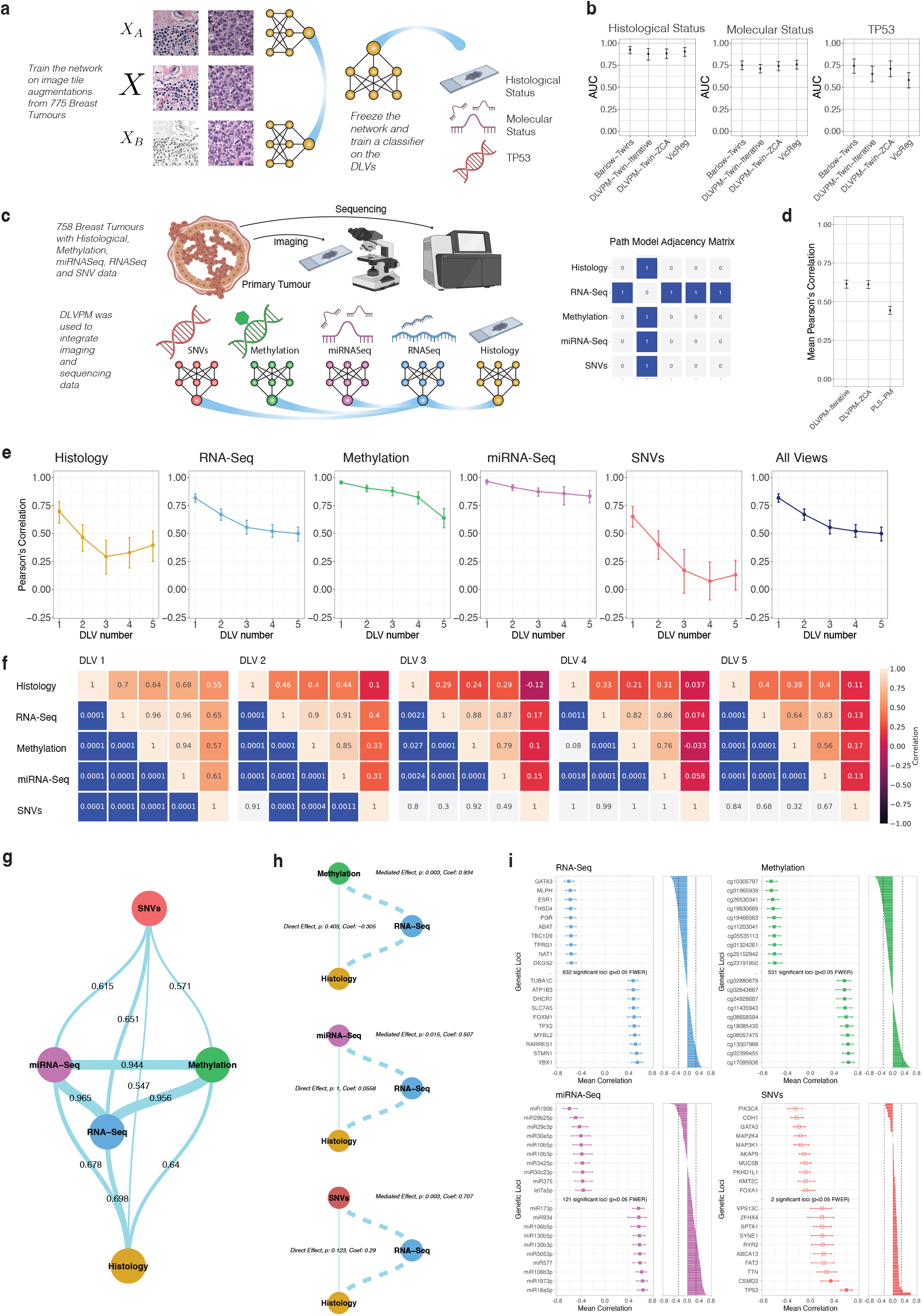
This figure illustrates the process by which DLVPM can be trained in a siamese/twin network configuration. b: These plots compare the performance of DLVPM-twins against the performance of VicReg and BarlowTwins. The error bars represent 95% confidence intervals. c: Conceptual illustration of the DLVPM method, and the multi-omic imaging data it is applied to here; this panel shows a graph representation of the path model, and the associated adjacency matrix. d: A comparison of the mean Pearson’s correlation across both orthogonal dimensions and data modalities for DLVPM using both ZCA-whitening, and iterative orthogonalization procedures, and the shallow PLS-PM method. e: For each data type, these plots show the mean Pearson’s correlation of each DLV, with DLVs from data types connected by the path model. The error bars on the plot denote 95% bootstrapped confidence intervals. f: Association matrices for all five DLVs. Entries in the upper triangular part of the matrix indicate Pearson’s correlation values between different data-types. Entries in the lower part of the matrix are significance values for these correlations, obtained using permutation testing. g: A path model linking the omics and imaging data types included in this analysis. This network graph represents the first orthogonal mode of variation between DLVs, established using the DLVPM procedure. The edges connecting network nodes are labelled with the Pearson’s pairwise correlation coefficient value calculated between DLVs. h: Results of mediation analyses carried out using the first DLV. The numbers on the network graph show beta values and significance levels for the direct and mediated effects. i: Results of additional analyses to localize effects to particular genetic loci. The plot shows Pearson’s correlation values between genetic loci, and DLVs connected to the data-view under analysis by the path model. The plots on the left shows the ten most positively and negatively associated genetic loci for each data type. The error bars represent 95% bootstrapped confidence intervals. The bar plots show Pearson’s correlation values for all loci under analysis, along with the familywise error corrected significance threshold. Due to space limitations, we only show analyses for the first DLV here.

Twin networks are typically benchmarked by comparing their performance on downstream tasks ^28^. We compared DLVPM-twins to the VicReg and BarlowTwins methods by training a single layer classification-head on top of the embeddings learned by each method. We applied DLVPM to data from 775 breast cancer samples from The Cancer Genome Atlas (TCGA). Each model was trained using the EfficientNetB0 architecture^29^. We used this classification head to predict: Histological and Molecular status, and the presence of TP53, the most common mutation in breast cancer^28,30^. Training and testing were carried out on a random 80% to 20% split of the data.

We found that the performance of both our formulations of DLVPM-Twins equalled the BarlowTwins and VicReg procedures (see Figure 2b). Our method has the advantage that it does not require any hyperparameters other than the size of the final embedding. Fewer hyperparameters simplify the training process and increase model robustness by reducing the need for extensive tuning. Among the two DLVPM variants, the version employing iterative orthogonalization has the major advantage that it ranks the latent variables according to the strength of their associations across data types, in a manner akin to the ranking of principal components on the basis of the variance they account for within a dataset. Further, DLVPM can be used to integrate data from arbitrarily many data-types. In contrast, Barlow Twins can only learn a representation of a single data type; VicReg can learn representations of two data types but it is unclear how the method can be generalized further than this.

### Full Deep Latent Variable Path Modelling

Next, we trained DLVPM for the purposes of full path modelling. We applied DLVPM to data from 758 breast cancer samples from The Cancer Genome Atlas (TCGA). Our initial goal was to uncover relationships across five types of data: histological images, single nucleotide variants (SNVs), methylation profiles, miRNA sequencing (miRNA-Seq), and RNA sequencing (RNA-Seq) expression data. We positioned transcriptomic data at the centre of the path modelling analysis, recognizing its pivotal role in mediating the effects of genomic and epigenomic changes on histological tissues through gene expression modulation. Using this path model, all other data types are linked to one another indirectly through the RNA-Seq data (see Figure 2c). Training and testing were carried out on a random 80% to 20% split of the data.

We must also specify measurement models for each data-type, these models define the manner in which the data is processed and connected. For the histology data, we specified a neural network designed to mimic the way that a histologist would work: The method aggregates effects arising at different magnifications, here it takes effects at x5, x10 and x20 magnification, each utilizing the EfficientNetB0 convolutional architecture^29^. For the genetic data, we used a residual network with an attentional mechanism. This allows the neural network to aggregate the linear effects from individual genes, with interaction effects between genes. The full neural network encompassing the path model and individual measurement models is shown in Extended Data Figure 3.

Following model training, we compared the performance of DLVPM to Projection to Latent Structures Path Modelling PLS-PM^19^. PLS-PM has an identical objective to DLVPM, but is only able to model linear effects. In this comparison, both the iterative orthogonalization and ZCA-whitening versions of DLVPM demonstrated greatly superior performance compared to PLS-PM (Figure 2d). As previously noted, the DLVPM variant utilizing iterative orthogonalization has the advantage that it ranks DLVs by their importance. For this reason, we used the results from this approach in all subsequent analyses. This ranking of DLVs by their mean association is shown in Figure 2e for both the model as a whole, and for each data-type individually.

Next, we evaluated the specific associations the method uncovers between data-types. These associations and their permutation family-wise error corrected significance levels are shown in Figure 2f. This analysis uncovers multiple orthogonal paths that connect molecular and histological data. A network graph illustrating the DLVPM path model for the first set of DLVs is illustrated in Figure 2g.

A major strength of DLVPM is its ability to uncover and analyse indirect effects, such as mediation relationships among variables, opening up the possibility of investigating the intricate dynamics and indirect effects that define complex systems. We examined how RNA-Seq DLVs mediate the interaction between various genetic and epigenetic variables—specifically Methylation, miRNA-Seq, and Single Nucleotide Variants (SNVs), which are treated as independent variables—and histological outcomes, which are treated as dependent variables. Path diagrams, presented in Figure 2h, visually depict these mediation processes, highlighting both direct effects and those mediated via the RNA-Seq DLV. Notably, the RNA-Seq DLV serves as a full mediator, meaning it entirely accounts for the relationship between the independent and dependent variables. This analysis highlights the crucial role of gene expression in linking genetic and epigenetic changes to cellular and tissue-level phenotypes, offering insights into the complex interactions that drive histological changes (See Figure 2h and Extended Data Figure 4).

Consistent with DLVPM’s path model, which links all data-types through the RNA-Seq data, we observed that all DLVs, even those originating from histology data, demonstrated a stronger association with established clinical molecular subtypes than with histological types. For instance, the first DLV distinctly stratified basal and luminal molecular subtypes across all data modalities, as shown in Extended Data Figure 5.

DLVPM operates fundamentally as a multivariate approach, designed to uncover factors exhibiting high correlation across diverse data types, including genetic and imaging datasets. The multi-omic DLVs constructed by the model represent complex polygenic factors. Due to its multivariate nature, the method initially precludes direct attribution of significance to specific genetic loci within the model. To bridge this gap, we ran additional analyses to isolate genetic/epigenetic loci that demonstrate significant correlations individually with DLVs (see Methods). Each DLV produces a stratification of imaging/multi-omic subtypes, with loci exhibiting either positive or negative associations. Hundreds of loci made individually significant contributions to the DLVPM model (Figure 2i, Extended Data Figure 6, Supplementary Table 1). Permutation testing using the distribution of the maximal statistic was used to control for multiple comparisons and provide strong control over the family-wise error rate.

The first DLV shows the strongest associative mode linking the omics and imaging data, and effects on histology are fully mediated via gene expression, quantified by RNA-Seq. This prompted us to focus our initial interpretation of individually significant loci on RNA-Seq data from DLV 1. First, investigating negatively associated loci: this path model stratifies genes important in luminal-basal transcriptional differentiation program: ESR1, which defines the luminal subtype, is used for clinical diagnosis of breast cancer, and is a target in hormone therapy ^31^. Further, GATA3, which exhibits the strongest negative association with DLV 1, regulates luminal cell differentiation and exhibits a shift from a tumour-suppressing to a tumour-promoting role in breast cancer via the deregulation of THSD4^32^, which also shows a strong negative association. MLPH emerges as having the second strongest negative association with DLV 1, and is also used diagnostically^33^. The PGR gene, which is crucial for prognosticating hormone treatment outcomes in breast cancer is also significant here, and is closely linked to luminal breast cancer^34^. In contrast: genes showing a strong positive association with DLV 1 have been primarily linked to the basal breast cancer subtype: YBX1 has been significantly implicated in breast cancer, and is particularly noted for its role in cell migration and invasion^35^, and drug resistance^36^. STMN1 has been implicated in cell cycle progression and mitosis and has been investigated as a therapeutic target in breast cancer^37^. RARRES1 is highly expressed in basal breast cancers^37,38^. MYBL2 has been shown to drive cell cycle progression in breast cancer^39^. A luminal to basal stratification on the first DLV was supported by gene set enrichment analyses carried out between DLVs and gene expression scores (see Methods); other DLVs were associated with different cancer related processes (Extended Data Figure 7).

### Characterization of Histological Data

A number of important previous studies have leveraged histological data to predict clinical molecular status, detect the presence or absence of known oncogenic mutations, and delineate bulk transcriptomic profiles of tumours using deep neural networks^11, 40^. While these studies are hugely important, they largely operate within the confines of pre-existing hypotheses. The DLVPM methodology stands out for its capacity to unearth previously unrecognized relationships across diverse data modalities.

We ran further analyses focused on pinpointing multi-omic loci that show individually significant correlations with histological DLVs, which represent an outcome phenotype. Our investigations not only corroborated existing knowledge by identifying histological-genetic associations with well-documented oncogenes such as TP53, but also uncovered significant links between histological features and hundreds of previously uncharted multi-omic loci (Figure 3a, Extended Data Figure 8, Supplementary Table 2).

**Figure 3a:**
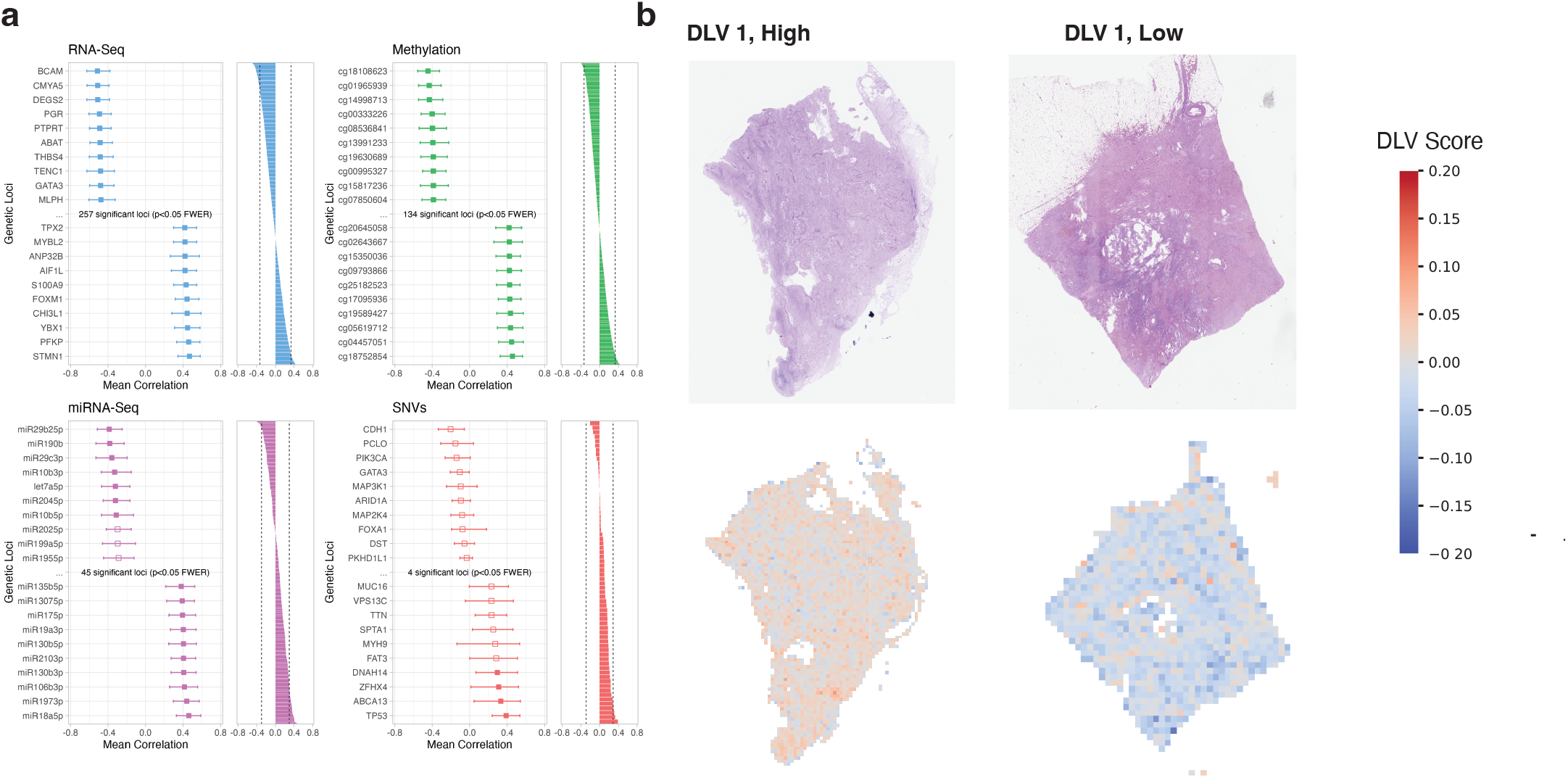
Results of additional analyses to localize effects to identify omics loci showing an individually significant association with histological data. The plot shows Pearson’s correlation values between genetic loci, and the first histological DLV. The plots on the left shows the ten most positively and negatively associated genetic loci for each data type. The error bars represent 95% bootstrapped confidence intervals. The bar plots show Pearson’s correlation values for all loci under analysis, along with the familywise error corrected significance threshold. Due to space limitations, we only show analyses for the first DLV here. b: Heatmaps for high scoring and low scoring tumours, on the first DLV, at x10 magnification.

The first histological DLV exhibited by far the largest number of individually significant associations with multi-omic loci (Figure 3a, Extended Data Figure 8, Supplementary Table 2). Many of the genes making the strongest individual contribution to the overall model were also amongst those showing the most pronounced correlations with the histology data: GATA3 and PGR both showed a strong negative association with DLV 1, both these genes have been reported as exhibiting pronounced inverse correlations with histological grade, mitotic rate and nuclear pleomorphism in hypothesis driven studies involving expert pathological assessment^41^. The cell adhesion gene BCAM, which has been previously linked to epithelial cancers^42^, shows the strongest negative association with DLV 1. Genes showing a strong positive association with histology are primarily associated with the more aggressive basal type. TPX2, ANP32B, PFKP, CHI3L1, S1009A, FOXM1, STMN1 and MYBL2 are all crucially involved in cellular proliferation^43,44,37,39,45–48^. This is particularly noteworthy in this context where increased cellular proliferation will result in visible changes in tumour grade and mitotic rate.

As was previously noted, we trained our model on small tissue sections known as image tiles (see methods). Once the DLVPM model is trained, we can deconvolve tile-wide effects back into image space. We used a novel neural network model that takes tiles at x5, x10, and x20 magnifications (see Extended Data Figure 3). Figure 3b shows high and low scoring tumours at x10 magnification, stratified on DLV 1. Histological effects and their molecular concomitants are explored in greater detail later in the text.

### Single Cell Characterisation

DLVPM acts to simultaneously integrate multi-omic and imaging data, and reduce its dimensionality. This results in a compressed representation of important genetic and physiologic processes in cancer into a small number of Deep Latent Variables (DLVs). Once the overall model is trained, the individual measurement sub-models can be used to help further characterize the model, and obtain deeper biological insights. For this purpose, we applied the trained DLVPM model to single-cell, cell-line and spatial transcriptomic data.

Cancer cells are the fundamental units of neoplastic disease. Tumours are composed of a diverse array of cancer and stromal cells, with distinct genetic and phenotypic properties. We applied the RNA-Seq component of the full DLVPM model, trained on TCGA, to data from the single cell breast cancer encyclopaedia, which contains RNA-Seq data from 100,064 single cells^49^. This allows us to determine individual cell types that contribute significantly to each DLV, providing increased phenotypic resolution and potential insight into heterotypic interactions between cells that are typical of tumours scoring highly on different DLVs.

In concordance with earlier results, the first DLV exhibits strong negative enrichment for luminal cells types; this DLV also shows strong positive enrichment for basal and cycling cancer cells, which indicate a more aggressive molecular type (Figure 4a). The third DLV stratifies cells by their HER2 status, which represents an intrinsically aggressive molecular type, but which is amenable to targeted therapy^50^.

**Figure 4a:**
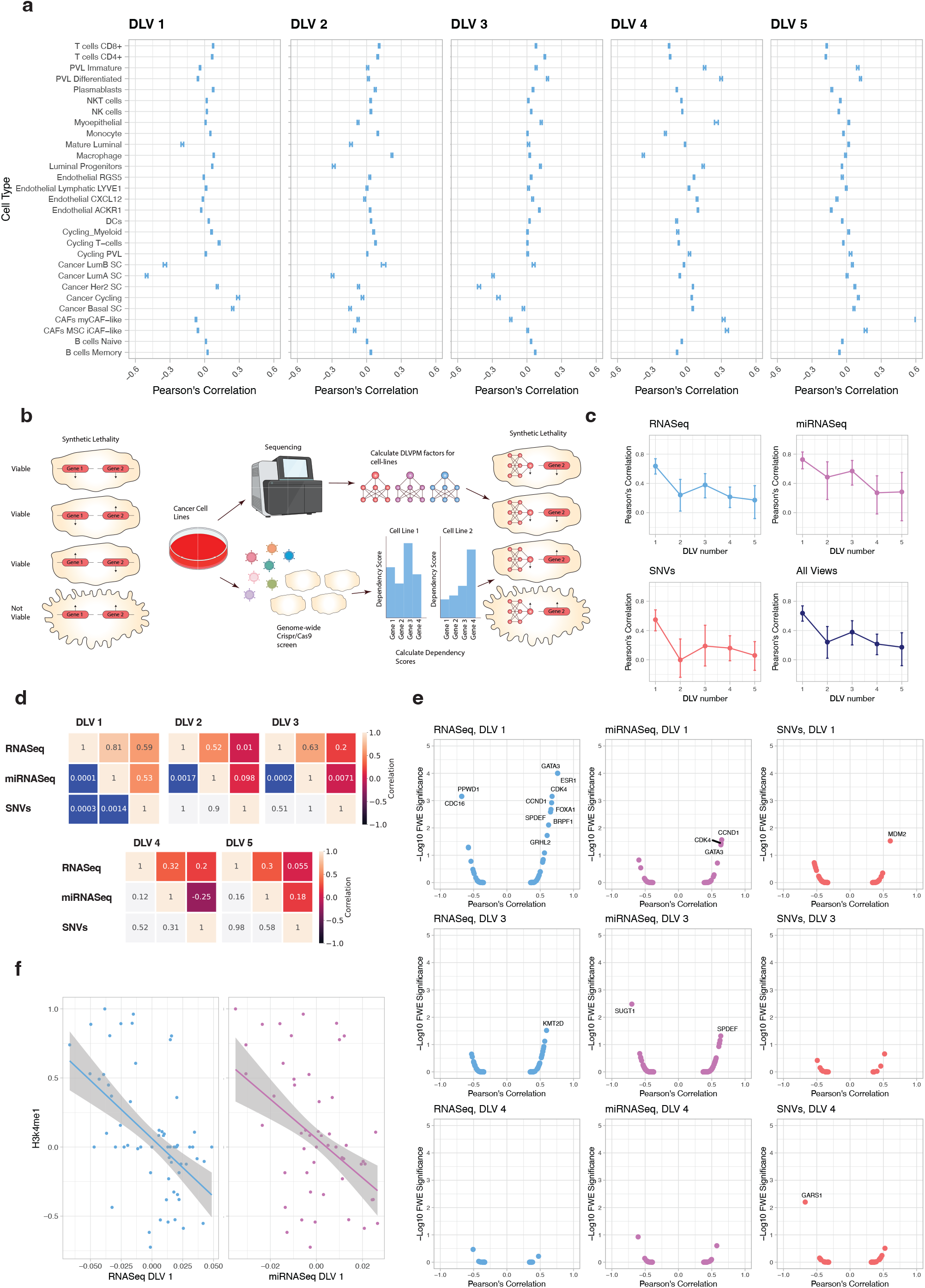
Shows the stratification of single cells based on the DLVPM model, applied to their transcriptomic profiles. The error bars show 99.9% bootstrapped confidence intervals. b: A conceptual illustration showing the general principle of synthetic lethality. Up and down arrows associated with the genes represent different states, for example: mutated/non-mutated. Genetic features are synthetic lethal if cell viability is affected when they both take on a particular state in the cell. Here, that is represented by up-arrows. The right part of the panel shows how DLVPM can be used to uncover new genetic vulnerabilities using this principle. DLVs are constructed from sequencing data, and are used to predict susceptibility to gene knockout. c: For each data type, these plots show the mean Pearson’s correlation of each DLV, with DLVs from data types connected by the path model. The error bars on the plot denote 95% bootstrapped confidence intervals. d: Association matrices for all five DLVs. Entries in the upper triangular part of the matrix indicate Pearson’s correlation values between different data-types. Entries in the lower part of the matrix are significance values for these correlations, obtained using permutation testing’s: Magnitude of associations between DLVs and CRISPR-Cas9 gene dependency scores, against their FWE corrected significance levels. Labelled genes are those with significance levels under p=0.05. We only show the volcano plots where there was a significant association between the DLVPM variable and the CRISPR-Cas9 data. f: Associations between the first RNA-Seq DLV and first miRNA-Seq DLV, and the histone modification H3K4me1.

Interestingly, DLV 5 shows extremely strong positive enrichment for myCAF fibroblasts, a subtype of CAFs identified by their expression of alpha-smooth muscle actin (α-SMA), which contributes to their effect on the tumour microenvironment, affecting tissue stiffness, cancer cell invasion and immune suppression, making them a significant marker for aggressive cancer types^51^. Results from this secondary analysis highlight the capability of the DLVPM model to elucidate complex cellular interactions within tumours, enhancing our understanding of cancer cell dynamics and tumour heterogeneity.

### Cancer Cell Line Characterisation

To investigate the model’s utility further, we applied the DLVPM model (trained on TCGA patient data) to multi-omic cell line data from the Cancer Cell Line Encyclopaedia (CCLE). Our objective was to explore if breast cancer cell lines, stratified on the basis of DLV profiles, exhibited differential sensitivity to genome-wide knockouts, facilitated by CRISPR-Cas9 loss-of-function screens. This approach not only promises to enhance our understanding of the biological significance of each DLV but also identifies potential therapeutic targets by pinpointing gene knockouts that exhibit synthetic lethal interactions in specific cancer contexts. This analysis utilises data from the cancer dependency map^52^. A schematic of the analysis is shown in Figure 4b.

We first tested if associations between omics DLVs, specified by the DLVPM model trained on TCGA patient data replicated in cell line data from the CCLE. We found that the first four DLVs retained significant associations (Figure 4c), with correlations between RNA-Seq and miRNA-Seq DLVs exhibiting a particularly large effect (Figure 4d).

We then conducted analyses to identify synthetic lethal interactions between DLVs, and genome-wide CRISPR-Cas9 dependency scores. Without using any biologically informed priors, and using a model trained on patient rather than cell line data, we identified several genes that are already targets of frontline therapies in breast cancer (Figure 4e). These associations were linked to DLV 1: As previously noted, ESR1 encodes an estrogen receptor and ligand activated transcription factor. The protein encoded by this gene regulates the transcription of many estrogen inducible genes involved in growth, metabolism, gestation and sexual development. Endocrine therapy to inhibit estrogen is an extremely important therapy for estrogen receptor positive breast cancers^31^. The first RNA-Seq DLV also shows a dependence on both the cyclin dependent kinase CDK4 and its regulator CCND1. Higher CCND1 expression has been linked to an increased risk of death in ER+ breast cancer^53^. Drugs designed to inhibit the action of CDK4 have recently been shown to improve prognosis in hormone dependent cancers, beyond the use of endocrine therapy alone^54,55^. GATA3 and FOXA1 showed very high synthetic-lethal dependency relations with DLV 1 in this cell line data. As previously noted, the GATA3 protein has been shown to be crucial in the development of the mammary gland, and is critical to the luminal cell program in the breast^56,57^. Both FOXA1 and GATA3 are pioneer factors, a special type of transcription factor that can bind directly to chromatin. Pioneer factors have been called the master regulators of the epigenome, and of cell fate^58^, which operate by opening previously inaccessible regulatory elements. These factors have the largest effect on transcription via histone modification and chromatin remodeling^58^. We calculated the association between RNA-Seq DLVs and cellular global chromatin profile data. The first RNA-Seq DLV was strongly associated with the histone modification h3k4me1 (see Figure 4f). It has been shown that FOXA1 recruits enzymes responsible for these epigenetic modifications^59^.

The SNV component of DLV 1 shows a strong dependency relation with MDM2. This DLV is largely driven by TP53 (Figure 4e); the MDM2 gene encodes a protein that negatively regulates the TP53 gene by targeting its protein product for degradation, thereby controlling cell cycle regulation and apoptosis. This interaction forms a feedback loop crucial for cellular balance, but its disruption can contribute to cancer progression, making MDM2 a target for therapies aiming to reactivate the tumour repressive functions of p53^60^.

Cells with low scores on DLV 1 were susceptible to knockout of a wholly different set of genes (Figure 4e). The strongest dependency relation was with CDC16; this gene is part of the APC complex, which governs exit from mitosis^61^. PPWD1 is thought to be involved in protein folding^62^, but its role in cancer is less well studied.

DLV 3 and DLV 4 exhibited different sets of dependency relations. The RNASeq component of DLV 3 shows a strong association with KMT2D, which plays a vital role in cell-differentiation in breast cancer^63^. SUGT1 is negatively associated with DLV 3; this gene has been previously linked to tumorigenesis in the breast^64^. SPDEF is a transcription factor that has been linked to worse prognoses in breast cancer^65^. The SNV component of DLV 4 shows a significant association with GARS1, which has been identified as a potential oncogene in breast cancer^66^.

We repeated these analyses using RNAi loss of function screen and obtained largely similar results (see Extended Data Figure 9). Following multiple comparison correction, no significant dependency relations were uncovered using individual genetic features, highlighting the benefit of our polygenic, compared to monogenic approaches. In summary, our cancer dependency map analysis underscores that it is possible to identify genes essential to the cellular functioning of certain molecular subtypes of breast cancer using a DLVPM model trained on patient data. Our findings suggest that DLVs can serve as biomarkers for identifying cell lines that are particularly susceptible or resistant to specific genetic interventions, underscoring the potential for DLVPM-guided targeted therapies in oncology.

### Spatial Transcriptomics

Our analyses using the DLVPM model identifies histological concomitants of polygenic modes of variation in cancer, and their synthetic lethal genetic dependencies. Spatial transcriptomic data offers a more spatially resolved view of the genetic basis of aberrant changes in tissue structure. We found the genes ESR1, GATA3, CCND1, and FOXA1 to be highly dependent on DLV1 based on gene dependency scores from CRISPR-Cas9 screens. Each of these genes showed an individually significant association with histology (Figure 3a, Extended Data Figure 8, Supplementary Table 2), and are also included as part of the Xenium spatial transcriptomic gene panel^67^. We used Xenium spatial transcriptomic data to investigate the association between expression of these genes, and the histological concomitants of DLV 1 across individual tumours. We assessed tile-wise effects at x20 magnification, as this resolution is closest to the subcellular Xenium resolution.

We found significant associations between the spatial distribution of each of these genes, and the histological component of DLV 1 (see Figure 5a and Extended Data Figure 10) in both invasive ductal carcinoma and invasive lobular carcinoma, the two most common histological types of breast cancer. The strong association between these genes is indicative of their close functional relationship in breast cancer. Each of the four genes is most highly expressed in relatively well differentiated tumoral regions. This is where the histological component of DLV 1 also scores lowest. This is consistent with the suggested role of these genes in the early stages of tumour growth and progression.

**Figure 5a:**
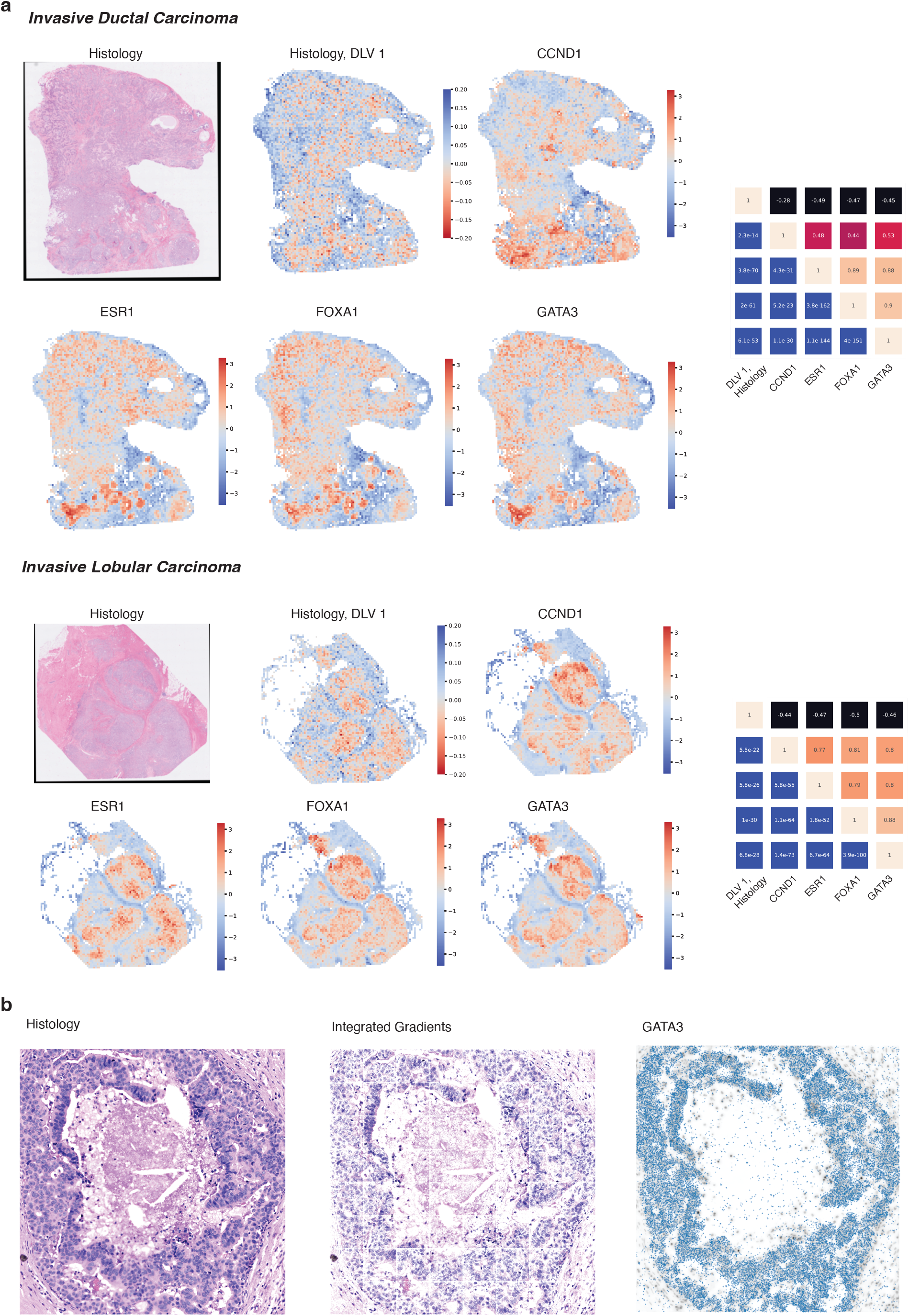
Tile-wise heatmaps generated from the DLVPM model, trained on TCGA data, and applied to histological and associated spatial transcriptomic data. The colormap is flipped for the histology heatmap as this DLV shows a negative association with the genes of interest. We applied this analysis to invasive ductal carcinoma and invasive lobular carcinoma. The association/significance matrices on the right show correlations between genes of interest and the first histology DLV for both tumours. The upper triangular part of each matrix is denoted with Pearson’s Correlation Coefficient between each gene, and the histology data. The lower triangular part of each matrix denotes the significance level between genes and histology data. b: The image on the left shows a cancerous breast duct, which scored highly on DLV 1. The middle panel shows a feature attribution map, generated using integrated gradients, illustrating the regions of the tumour that contributed most to the DLVPM model. The image on the right shows spatially mapped GATA3 transcripts.

In order to pinpoint histological features that play a pivotal role in linking genetic profiles with histological patterns at a more granular, sub-tile resolution, we applied the Integrated Gradients method for feature attribution^68^ (see Figure 5b). This attribution technique assigns significance scores to specific image regions, thereby identifying those that have a pronounced influence on the predictions made by DLV 1. Of particular interest, well differentiated ductal regions received elevated scores, highlighting their marked importance in the model’s determination. Furthermore, these regions show a high concentration of key genetic markers, including GATA3, CCND1, FOXA1, and ESR1. The DLVPM analyses carried out on this Xenium data forge a connection between the functional essentiality of genes, as assessed by CRISPR-Cas9 loss-of-function screens, and their spatial expression patterns, framing an integrated model of disease pathology.

## Discussion

DLVPM is a new method for modelling dependencies between different data types. This method stands out for its ability to uncover complex, non-linear interactions among both structured and unstructured data-types, overcoming the limitations of traditional path modelling techniques^19^. Initially trained on the extensive TCGA dataset, the modular nature of the method allows for flexible adaptation and further refinement with additional datasets such as single-cell, cell-line and spatial transcriptomic analyses. By applying DLVPM to the cancer dependency map data, it unveils critical insights into multi-omic dependencies. Furthermore, DLVPM bridges microscopic tissue structure changes with genetic vulnerabilities identified by the same model, illustrating its ability to construct holistic models of illness pathology. This method’s comprehensive data integration capability marks a significant step forward, promising applicability beyond cancer to a broad spectrum of diseases.

DLVPM is superior to classical approaches to path modelling in terms of the magnitude of associations the method is able to establish between different data types. This is likely because, in contrast to classical approaches, this method is able to model the complex breakdown in molecular machinery that underpins carcinogenesis: Typically, cancer initiation requires mutations or epigenetic changes in several driver genes. These alterations can affect gene expression at thousands of loci. Many of these genes will be transcription factors, whose purpose is to control the expression of other genes. Classical methods are unable to parse this complexity as they are only able to model linear effects. In contrast, deep learning methods have already been shown to be capable of modelling complex interactions between loci across the genome^12,13^.

Historically, researchers have developed drugs that target oncogenes and block their function. However, not all cancers have oncogenes, which limits the number of possible drug targets. To overcome this challenge, researchers have utilised the principle of synthetic lethality^69^. Synthetic lethal interactions in cancer can be probed using functional genetic experiments such as CRISPR-Cas9 loss of function screens. Associations between cells’ molecular features, and susceptibility to knockout of a particular gene, represents a synthetic lethal interaction. This approach also fails to respect the intrinsically polygenic nature of cancer as an illness. DLVPM simultaneously integrates the multi-omic data it is applied to, and reduces its dimensionality, resulting in a small number of polygenic, multi-omic DLVs, avoiding both major pitfalls associated with taking a single gene approach.

DLVPM is implemented in the flexible and user-friendly TensorFlow/Keras ecosystem, enabling the modular construction of complex models tailored to a wide array of data analysis tasks. Using pre-defined Keras layers, users can define new DLVPM models in just a few lines of code. This modular design not only simplifies the development and testing of sophisticated models but also enhances their extensibility, ensuring that our method can be seamlessly applied across diverse research fields and data types. This toolbox also contains a sub-module for confound removal, which can also be used in classification and regression problems, and we anticipate as being generally useful to the Deep Learning field.

Many illnesses arise as a result of complex interactions between multiple biological and environmental factors. Several, large, open access databases, such as UK Biobank^70^, the European Genome-Phenome Archive (EGA)^71^, and the Cancer Imaging Archive^72^ have been created to help understand these factors, and contain large amounts of multi-omics and imaging data. Further, new technologies such as single-cell^73^ and spatial multi-omics^74^ produce enormous quantities of data that need to be integrated and reduced to be understood. DLVPM is ideally suited for this task, as it is able to link arbitrarily many data modalities, including both structured and unstructured data. In this investigation, we showed how DLVPM can be used to construct a global model of breast cancer as an illness. Using this method, wholly different but connected manifestations of the same underlying illness can be understood with reference to the same neural network model.

## Methods

### Deep Latent Variable Path Modelling

In the text that follows, we give a general introduction and full technical treatment of the DLVPM method. DLVPM can be thought of as a generalization of projection to latent structures path modelling (PLS-PM)^19^. PLS-PM can be considered, in turn, to be a generalization of canonical correlation^75^. It is therefore natural to build an understanding of DLVPM with reference to these simpler methods. It is worth noting that we could have called our method Deep PLS-PM. However, we felt that DLVPM was more descriptive. We also wished to avoid confusion with the more popular PLS regression procedure.

The description of the DLVPM method we present here is broken into three basic parts:

1. A description of shallow (i.e. non deep-learning based) methods for establishing correlations between different data types.
2. Deep neural networks and notation.
3. A description of DLVPM, and how deep learning can be used to identify complex, non-linear associations between different data-types.

#### Canonical Correlation Analysis

Canonical correlation analysis (CCA) is a statistical method used to identify linear relationships between two or more sets of variables^75^. This method can be thought of as a generalization of linear least squares regression. The objective of CCA is to identify a relationship between two (or more) sets of variables, where there is no distinction between which variables are considered dependent, and which are considered independent. This method identifies weights for each variable, such that the weighted sum of variables in each set is maximally correlated with the weighted sum of variables from the opposite set, assuming a linear relationship^75^.

Consider two matrices *X*_1_and *X*_2_, where each row denotes one of *n* observations, and each column denotes *p*_*1*_ *or p*_*2*_ features for *X*_1_and *X*_2_ respectively. CCA is optimized to find weight vectors *w*_1_ and *w*_2_ that maximise the association:

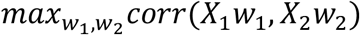

We assume that the columns of *X*_1_ and *X*_2_ have been standardised to have a mean of zero and a standard deviation of one. Using the equation used to find Pearson’s correlation coefficient:

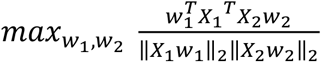

Notice that the denominator is simply a normalization term. Therefore, the canonical correlation objective can also be written as:

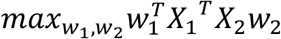

Subject to the constraints:

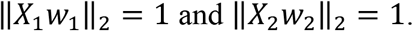

Where the vectors *X*_1_*w*_1_ and *X*_2_*w*_2_ are referred to as canonical variates.

In the original formulation, the canonical weights that maximize the association between the two data-views are normally found using eigenvalue decomposition. It is possible to find multiple modes of variation using this method. Here, the correlation between subsequent canonical variates is maximised subject to their being uncorrelated with other canonical variates. A total of *n*_*dims*_= *min*(*p*_1_, *p*_2_) canonical variates can be extracted in this way. This can be written as:

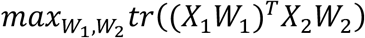

Subject to the orthogonalization constraint:

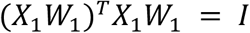

And:

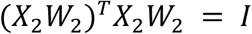

Where *tr* represents the matrix trace, *W*_1_ and *W*_2_ are *p*_1_ × *n*_*dims*_ and *p*_2_ × *n*_*dims*_ matrices respectively, and *I* is a *n*_*dims*_× *n*_*dims*_identity matrix.

#### Generalizing CCA

Hotelling’s original formulation of CCA was designed to identify associations between two data-views. Researchers have generalized CCA to more than two data-views^76^. There are a number of different ways in which this can be done. One way to generalize the CCA approach is to optimize the sum of correlations between different data-views. This involves maximizing the following criteria:

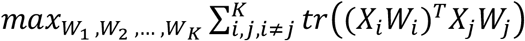

Subject to the orthogonalization constraints:

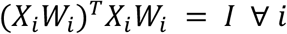

Where different data-types are denoted by *i*, and *K* represents the total number of data-types.

#### Projection to Latent Structures Path Modelling

The canonical correlation procedures described in the text above can be used to identify latent variables that are highly correlated between multiple data types. In some cases, we may wish to identify associations between some, but not all, data types. For example, a particular disease phenotype may have both genetic and environmental causes. It does not make much sense to try to link these genetic and environmental causes as they should be independent. Any model that attempts to link these data types may end up highlighting spurious effects.

The mathematical framework above, described with relation to generalised canonical correlation analysis, can be used to formulate a kind of structural equation modelling procedure called projection to latent structures path modelling (PLS-PM)^77,78^. Using PLS-PM, it is possible to identify associations between pre-specified data-types. Utilizing this method, we specify which data-types are connected with one another using a pre-defined Adjacency matrix *C*. The adjacency matrix is a square matrix where the elements *c*_*ij*_ represent connections between views *i* and *j*.

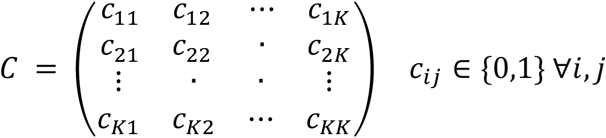

Where *K* is the total number of data-types under analysis.

The optimisation criteria can then be written as:

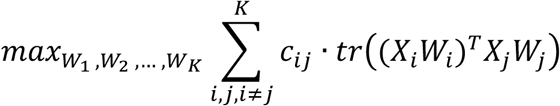

Subject to the constraints:

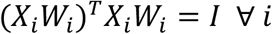

Where *c*_*ij*_ represents binary indexed elements of the adjacency matrix *C*. Using PLS-PM, the full modelling process is normally referred to in two parts: the structural model, and the measurement model. The structural model is the part of the model that defines which interrelations to optimize between data types, this information is stored in the adjacency matrix: *C*. The measurement model is the part of each model is denoted by *X*_*i*_ *W*_*i*_ ∀ *i*, this is the part of the model linking individual features to latent variables, in the path model^19^.

#### Deep Neural Networks and Notation

Neural networks are computational models composed of layers of interconnected “neurons” that perform calculations. During training, these networks adjust neuron connection weights via backpropagation, where they compute the gradient of a loss function (the difference between predicted and actual data) and iteratively update network weights. The outputs of most neural networks can be written in the very general form:

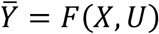

Where 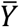 is the network output and *F*(*X, U*) is some function that takes an input *X* and passes it though sets of weights and biases *U*. This could be many kinds of neural network, for example a feedforward neural network, a convolutional network or a transformer. The network output can be written more simply still as: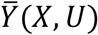.

Each of the methods described in the text below relies on the last layer of the neural network having a linear projection weight on the last layer. Treating this weight differently in the notation is crucial to understand the mechanisms by which DLVPM functions. Therefore, neural networks processing individual data-types in DLVPM are written as: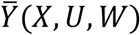 where *U* represents all weights and biases in the network up to the penultimate layer, and *W* represents the weights on the last layer of the network. We use this very simple notation to denote neural networks throughout the rest of the text.

#### Deep Canonical Correlation

Andrew et al developed a two-view form of canonical correlation analysis, which they termed Deep canonical correlation analysis (Deep CCA)^79^. Deep CCA creates highly correlated representations of two data types by passing them through deep neural networks. The goal of the algorithm is to learn weights and biases of both data-views such that we seek to maximise:

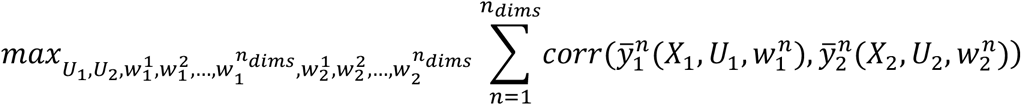

Subject to the orthogonalization constraint:

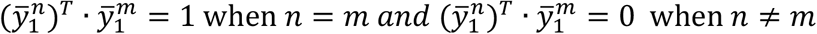

Where *n*_*dims*_ is the total number of canonical variates we wish to extract.

This optimization problem can be written in the matrix form as:

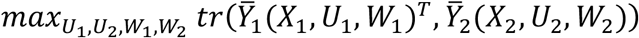

subject to the orthogonalization constraint:

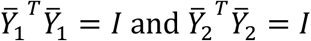

Where *tr* is the matrix trace, 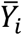 is a column-wise concatenation of 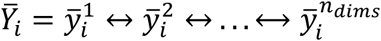, where ⟷ signifies the column-wise concatenation of CCA factors, and *I* is the identity matrix. Here, *W*_1_ And *W*_2_ represent the set of all weights: 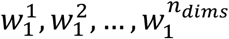 and 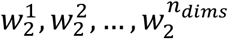.

Andrew’s formulation of this procedure operates by taking the derivative of the cross-covariance matrix between data-views. However, this approach is difficult to generalise to more than two data-views.

Wang et al^80^ formulated an iterative least squares approach to this method. This involves minimising the loss:

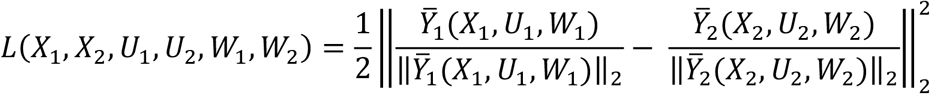

Subject to the orthogonalization constraint given above. We use a similar iterative least squares regression approach in the present investigation.

#### Deep Latent Variable Path Modelling

The goal of the DLVPM algorithm is to identify orthogonal modes of association between data-views connected by the user-defined adjacency matrix: *C*. As before, the adjacency matrix is a square matrix where the elements *c*_*ij*_ represent connections between views *i* and *j*. This method is essentially a deep analogue of PLS path modelling. This adjacency matrix is often referred to as the structural or path model.

We therefore seek to maximise:

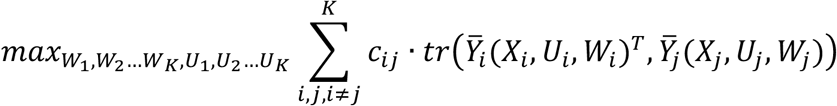

Subject to the orthogonalization constraint:

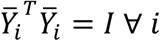

Taking the iterative regression approach followed by Wang et al, and described with reference to classical canonical correlation and PLS-PM earlier in the text, we can maximize the association between network outputs by minimizing the loss:

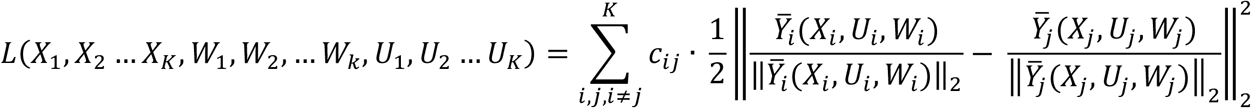

#### Orthogonalization

The DLVPM algorithm can be split into two fundamental parts: An optimization step aimed at finding factors that are strongly correlated between data-views, and a constraint that ensures that DLVPM factors are orthogonal to one another. It is possible to identify a single factor of shared variance between sets without the orthogonalization step. The loss associated with finding a single DLVPM factor can be written as:

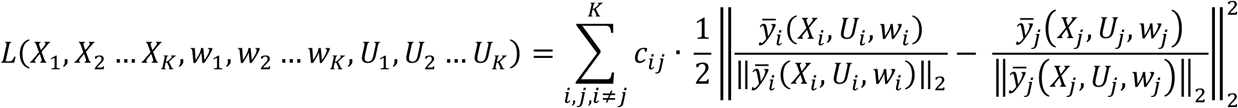

In cases where we wish to identify more than a single factor of shared variance between data-views, an orthogonalization step is required to de-correlate factors. We used two different approaches to orthogonalization in the present investigation. We first introduce an orthogonalization procedure inspired by classical PLS, which is used in the main part of this investigation. We also compared this approach to a whitening procedure, similar to the approach used by Wang et al^80^.

#### Iterative Orthogonalization

In the present investigation, we use a matrix deflation approach inspired by classical PLS-PM. This approach has the advantage that it maintains the proper ordering of DLVPM factors. During the forward pass through the network, data is orthogonalized with respect to previous DLVPM factors. Individual DLVPM factors are written as:

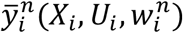

The set of all DLVPM factors in a data-view can be written as:

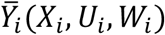

Where 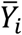 is an *N* × *n*_*dims*_matrix of DLVPM factors and *W*_*i*_ is a matrix of DLVPM weights. 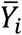 is a column-wise concatenation of 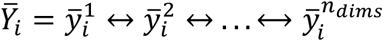. Similarly, we define the matrix 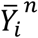 as the concatenation of all vectors from the first to the n^th^.

We denote the penultimate layer of the neural network with the notation: *F*_*i*_(*X*_*i*_, *U*_*i*_). It is a well-known property of regression that the residual features, denoted by 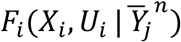, found in the regression:

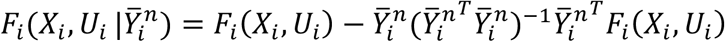

Are orthogonal to 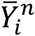 (utilizing the Moore-Penrose pseudo-inverse). We use this mechanism to identify orthogonal modes of variation using DLVPM.

We can then write the loss for the n^th^ extracted latent factor, for the i^th^ data-view as:

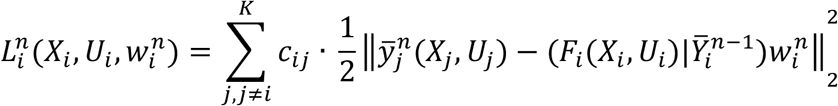

Given that 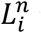 is a sum of regression problems. We can then write the total loss for the i^th^ data-view as:

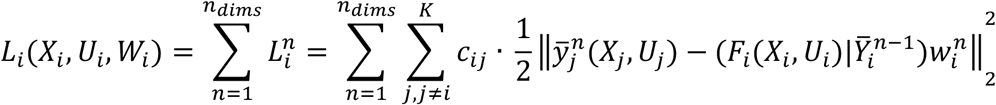

*L*_*i*_ is therefore written as a sum of mean squared error losses across latent factors. Similarly, the total loss can be calculated as the sum of losses across all data-views:

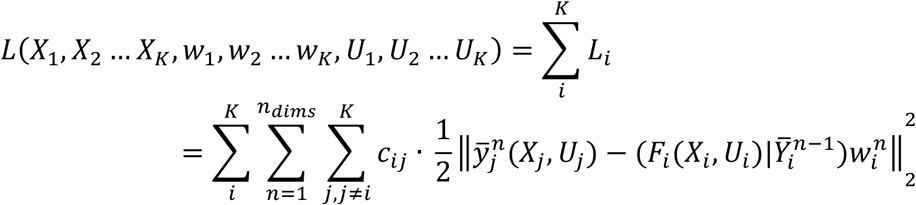

Due to the orthogonalization process introduced in the text above, this formulation meets the constraints:

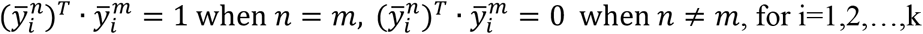

It is worth noting at this point that due to these constraints:

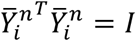

Where *I* is the identity matrix. This means that the orthogonalisation procedure:

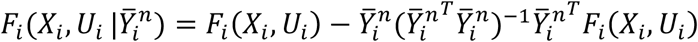

Simplifies to:

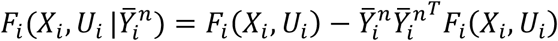

DLVPM minimizes this loss in an iterative fashion by calculating the gradients associated with each data-view and updating the weights of these data-views.

So far, analysis of the DLVPM algorithm has preceded assuming that training is carried out on the entire dataset simultaneously. However, neural networks are usually trained on subsets or batches of data while orthogonality is a property of the full dataset. Orthogonalization requires estimating the covariance matrix:

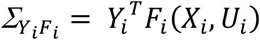

We calculate the covariance matrices above during model training, by making an initial estimate of the covariance matrices using the first batch, then updating this estimate using parameter re-estimation with momentum for each batch. The batch-level covariance matrices for the first batch are written as:

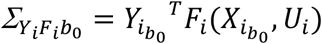

The global covariance matrices for the first batch are then initially estimated as:

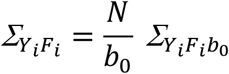

Where *N* is the total number of samples and *b*_0_ is the size of the first batch. In subsequent batch updates, covariance matrices are calculated as:

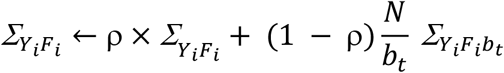

Where ρ is the momentum of the update, *N* is the total number of samples under analysis, and *b*_*t*_ is the size of the current batch. The momentum ρ is a hyper-parameter that defines how quickly the covariance matrices are updated using the current batch.

This algorithm allows us to learn global matrices for orthogonalization, which can then be used during inference. Nevertheless, we found that using these covariance matrices during training were ineffective at enforcing orthogonality. This is likely to be because using global covariance matrices does not enforce orthogonality at the batch-wise level. This means that gradient-updates can also be non-orthogonal. However, if we use batch-wise orthogonalization, this condition is strongly enforced.

Consider the sub-loss for a particular data-view, for a particular dimension of shared-variance:

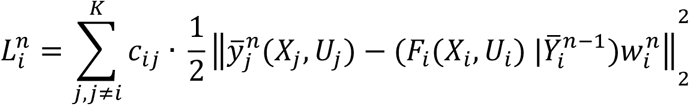

Taking the gradient of 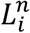 with respect to the weight 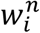 gives:

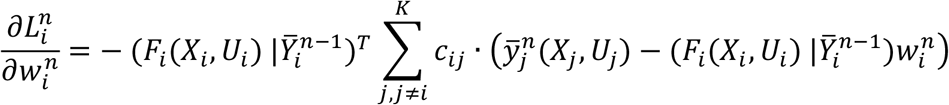

Using batch-wise covariance matrices:

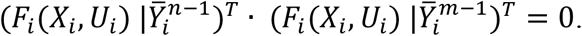

Therefore, gradients are orthogonal:

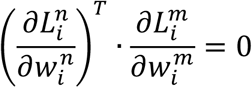

For these reasons, the algorithm we used functions differently in training and testing. During model training, we implement orthogonalization using batch-wise covariance matrices. Global covariance matrices are used during testing. This different behaviour during training and testing, using batch-wise and global parameters respectively, is similar in purpose and implementation to the batch normalization layer. The full algorithm specifying this method is shown in Figure 1 and Extended Data Figure 1. Pseudo code illustrating how this algorithm works is shown below:

#### Deep LVPM with Iterative Orthogonalization

**Input:** Data matrices 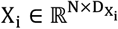 for i = 1,2, …, k. Initialization of the weights W_*i*_, U_*i*_ for each data-view, momentum p, learning rate 17. Randomly choose a mini-batch and extract data for each data-view: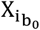.

**During Training:**

**For** t = 1,2,…,T, do:

Forward propagate through the network:

**For** i=0,1,2,…,*n*_views_, do:

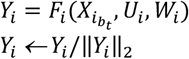

**For** i=0,1,2,…,*n*_views_, do:

Compute the batch mean and variance:

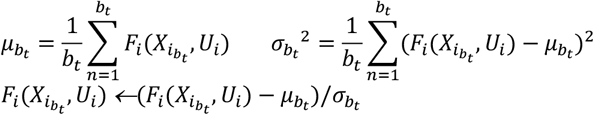

**For** n=0,1,2,…,*n*_dims_, do:

**If** n=0:

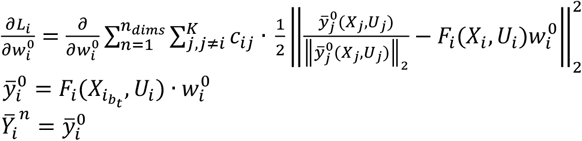

**Else:**

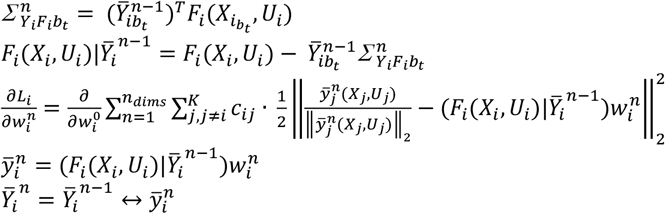

**If** t=0:

Define global variables, moving-mean and moving variance:

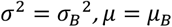

Covariance matrices (for orthogonalization):

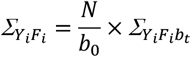

**Else:**

For subsequent batches, update the batch moving mean and moving variance:

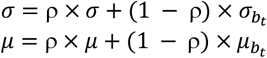

Update the moving covariance matrices:

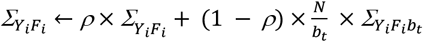

Update the weights:

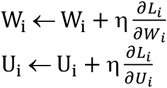

**During Inference:**

**For** i=0,1,2,…,*n*_v*i*ews,_ do:

Forward propaga*t*e through the ne*t*work:

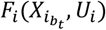

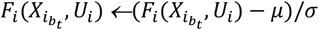

**If** n=0:

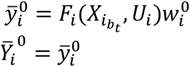

**Else:**

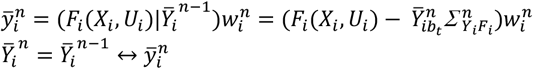

#### ZCA Whitening

ZCA whitening offers a different way of orthogonalizing deep latent variables. This method of orthogonalization was used by Wang et al in their formulation of deep canonical correlation analysis^80^.

Using the definitions of *Y*_*i*_ and *W*_*i*_ outlined earlier in the text. We can again write the objective as:

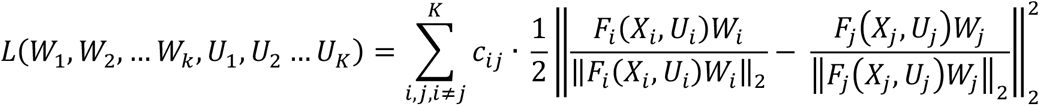

Subject to:

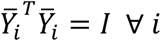

We note that if we multiply *Y*_*i*_ by the matrix square root of its inverse:

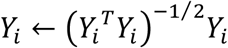

Then the columns of *Y*_*i*_, representing different modes of variation, are uncorrelated. In other words, the orthogonality condition is met.

We introduce a new algorithm, which again minimizes the global loss by iteratively minimizing the sum of squared loss between each data-view and connected data-views. Using the ZCA whitening approach, we iteratively minimize the loss:

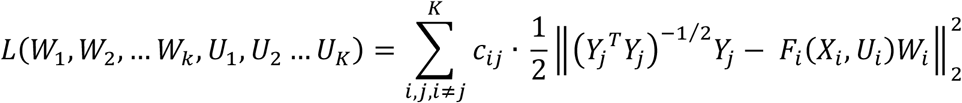

As was the case when minimizing the loss using the iterative orthogonalization approach specified above, when trained at the batch-wise level, we must estimate a global covariance matrix. We do this in a similar manner to the way in which we estimated global covariance matrices using the iterative orthogonalization approach. As noted in the explanation of the iterative orthogonalization algorithm, training using deep learning is generally carried out at the batch-wise level.

For the first batch, in each data-view, the covariance matrix is initially estimated as:

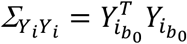

Subsequent batches are then estimated as:

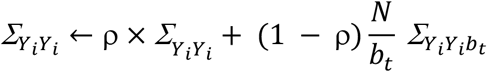

Full pseudo code for estimating a DLVPM model using the ZCA whitening orthogonalization approach is given below.

### Deep LVPM with ZCA whitening

**Input:** Data matrices 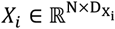, *k*.Initialization of the weights *W*_*i*_, *U*_*i*_ for each data-view, momentum ρ, learning rate η. Randomly choose a mini-batch and extract data for each data-view: 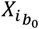.

### During Training

**For** t = 1,2,…,T, do:

Forward propagate through the network:

**For** i=0,1,2,…,*n*_views_, do:

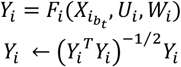

**For** i=1,2,…,*n*_views_, do:

Compute the batch mean and variance:

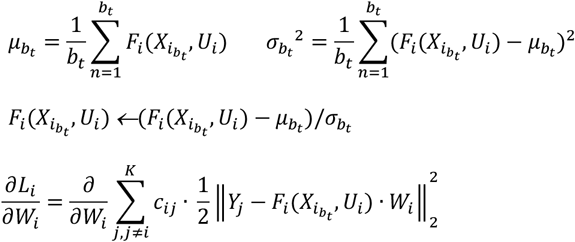

**If** t=0:

Define global variables, moving-mean and moving variance:

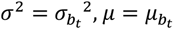

Covariance matrices (for orthogonalization):

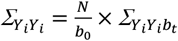

**Else:**

For subsequent batches, update the batch moving mean and moving variance:

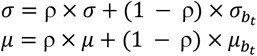

Update the moving covariance matrices:

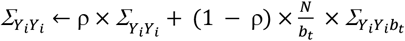

Update the weights:

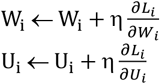

**During Inference:**

**For** i=1,2,…,*n*_views_, do:

Forward propagate through the network:

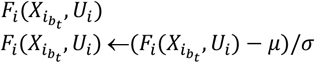

### DLVPM Twins

Although DLVPM is primarily designed to identify associations between multiple data-views, It can also be used to find useful representations of a single data-view, which can then be used for downstream tasks. When used in this manner, DLVPM falls into the class of methods called siamese or twin networks. This class of methods have become popular across a wide range of fields in recent years.

Twin architectures are trained by feeding a neural network distorted versions of the same input. By using some kind of correlative loss on the output features (as is the case with DLVPM), the network is encouraged to learn representations that are invariant to these distortions, which are likely to be useful in downstream analyses.

When using DLVPM in this way, the loss can be written as:

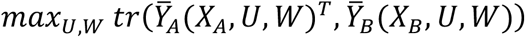

Where *tr* is the matrix trace, both subject to the orthogonalization constraint:

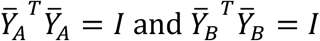

Here, *X*_*A*_ and *X*_*B*_ are augmented/distorted versions of the same input *X*. Note here that the weights and biases associated with the networks are the same for the entities we are seeking to maximise associations between. The network is then optimized to learn model outputs that are invariant to user specified changes in the input.

DLVPM-Twins can be used with both the iterative orthogonalization and ZCA-whitening approaches. The algorithms defining these approaches is very similar to the full path modelling algorithms.

Therefore, they are not given here in order to avoid repetition.

### Selecting Hyperparameters

Making the correct hyper-parameter selection choices is very important in order for a neural network to train smoothly. We found that a number of different hyper-parameter choices were important for proper training of the DLVPM model.

#### Learning Rate

The learning rate of a deep neural network determines the magnitude of each gradient learning update during model training. In general, using a high learning rate can lead to a deep neural network algorithm identifying a sub-optimal solution. When training DLVPM models, problems can also arise in the orthogonalization of DLVs. This is because increasing the learning rate causes an increase in the rate of change of network weights, and in turn, DLVs. It is then possible for changes in DLVs to occur too rapidly for an appropriate update of the covariance matrices used for orthogonalization. We used a learning rate of 1×10^−5^.

#### Batch Size

As with other correlation-based methods ^28^, we found that using a larger batch size tends to produce more stable results. In the current investigation, we used batch sizes of 256, which gave stable convergence.

#### Momentum

Momentum is a key parameter that must be specified for the update of covariance matrices using the DLVPM algorithm. This parameter is used to help get a stable representation of the covariance matrices linking DLVs, and the inputs to the DLVPM layer. If the momentum is set too low, the covariance matrix only reflects covariance in the most recent batches. If the momentum is set too high, the covariance matrix will not reflect recent updates very well. We found that a momentum value of ρ=0.95 works well when used for updating covariance matrices.

#### Dropout

Dropout helps to prevent overfitting. Overfitting is a particular problem when training multi-view methods such as DLVPM models. This is because the models are effectively trained to predict the output of one another. We used heavy dropout for each DLVPM measurement model.

#### Regularisation

Similar to dropout, L_1_ and L_2_-norm regularisation can help to prevent overfitting. We applied L_1_ and L_2_-norm regularisation to each of the DLVPM measurement models.

### Removing Confounds

Data is often subject to unwanted confounds, these confounds can affect the validity and generalisability of inferences made on this data. When assessing linear effects, these confounds can be removed by including them as covariates of no interest in a general linear model, or pre-regressing these unwanted effects from data prior to analysis. We took a similar approach to removing confounding effects in neural networks. The last layer of a DLVPM model is linear. Therefore, removing confound contributions prior to this layer will remove them entirely.

Here, a set of confounds is denoted by an N × D_c_ matrix *C*, where *N* is the number of samples, and D_c_ is the number of confounds. *F*(*X, U*) has the same definition, given earlier in the text.

Implementing the operation:

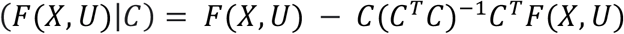

The matrix (*C*^*T*^*C*)^−1^*C*^*T*^ is known as the Moore-Penrose pseudo-inverse. It is a well-known result that columns of the resulting matrix: (*F*(*X, U*)|*C*), are orthogonal to the columns of the matrix *C*.

When using DLVPM, we can use this approach to orthogonalize neuronal outputs with respect to a set of confounds in the penultimate layer. As projection layers are linear, the outputs of the measurement model will be orthogonal to these confounds. As with the DLV orthogonalization described earlier in the documentation, we must adapt this orthogonalization so that it is possible to train models using this approach at the batch-wise level.

The matrix:

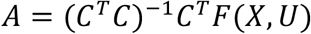

Can be split into two covariance matrices:

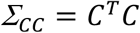

And:

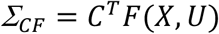

Batch-wise estimates of these matrices can be used to estimate full sample matrices. Batch-wise covariance matrices are written as:

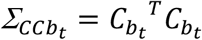

And:

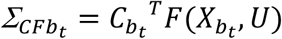

We can then carry out orthogonalization at the batch-wise level using:

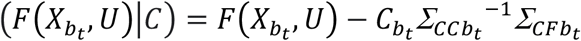

As in the case of carrying out orthogonalization between DLVs, we must also estimate full-sample covariance matrices so that we can carry out orthogonalization with respect to these parameters in unseen test data.

Global covariance matrices for the first batch are estimated as:

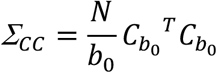

And:

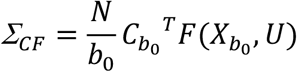

In subsequent batches, these covariance matrices are updated with momentum:

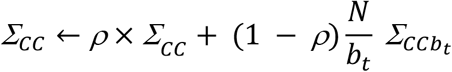

And:

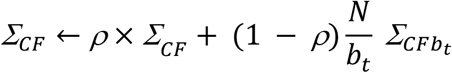

Where ρ denotes the momentum.

At model test time, these covariance matrices are then used to orthogonalize signal that is forward propagated through the network, with respect to confounding variables:

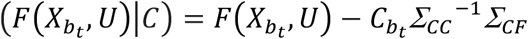

Full pseudo-code illustrating this process is given below.

**Confound Removal**

**Input:** Data matrices 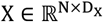, and confound matrices 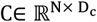.

**During Training:**

**For** t = 1,2,…,T, do:

Compute the batch mean and variance:

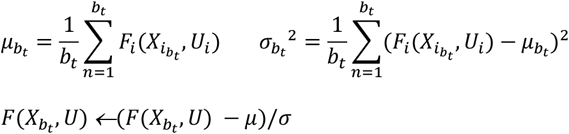

**If** t=0:

Define global variables, moving-mean and moving variance:

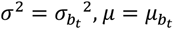

Covariance matrices (for orthogonalization):

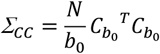

And:

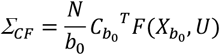

**Else:**

For subsequent batches, update the batch moving mean and moving variance:

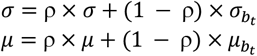

Update the moving covariance matrices:

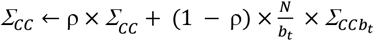

And:

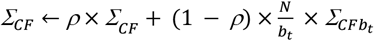

Use these covariance matrices to remove confounding effects:

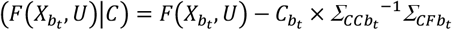

**During Inference:**

**For** i=1,2,…,*n*_views_, do:

Forward propagate through the network:

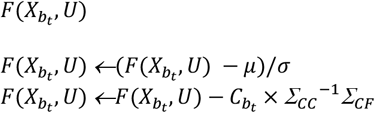

### Neuronal architecture

We must specify neural networks for each of the data types connected using the DLVPM algorithm. Different data types require the use of different neuronal architectures for processing and analysis.

The histological imaging data was processed using a network that aggregates effects visible in slide data at different magnifications. This is similar to the way in which a histologist will look for particular features at different magnifications. Tiles extracted at different magnifications were passed through an EfficientNetB0 architecture^29^, pretrained on the ImageNet dataset, to process effects at each of these magnifications. Global average pooling was carried out for each magnification. A feed forward neural network was then used to combine these multi magnification effects. The network utilizes skip connections from earlier to later layers of the network so that forward and backward signal propagation is unimpeded across the whole network. L_1_ and L_2_ weight regularization, along with a dropout layer^81^, was used to help prevent overfitting.

Each of the omics models uses the same general neural network structure. The model utilizes an embedding layer that reduces the dimensionality of the input to the square root of the initial gene count, a heuristic inspired by natural language processing to efficiently capture the essence of gene expression patterns. Subsequent reshaping introduces a pseudo-sequence dimension, enabling the application of a self-attention layer, which facilitates the model’s focus on critical gene interactions. The attention output, merged with the original input through a residual connection, ensures the preservation of initial gene expression information while incorporating learned interaction effects. Regularization was applied via L_1_ and L_2_ norms, scaled according to the dataset size, to mitigate overfitting, alongside a dropout layer for enhanced generalization.

Both the histological neural network and the gene data neural networks end with a custom neural network layer that partials out the effect of confounds using the Moore-Penrose pseudo inverse. This approach is detailed earlier in the Methods, and is illustrated in Extended Data Figure 3.

### Shallow PLS

We ran additional analyses to determine whether DLVPM is more effective than classical shallow methods when seeking to identify associations between different data types. There are two major types of PLS algorithm: mode A and mode B^19^. Mode A involves optimizing the association between different data types. This approach requires the calculation of the matrix inverse of within-modality covariance matrices. This is not possible when the number of examples in the data modality is smaller than the number of features. Mode B PLS solves this issue by replacing within-modality covariance matrices with identity matrices. As the data in the present application have many more features than samples, we used mode B PLS for comparison to DLVPM. Using shallow PLS, an EfficientNetB0 architecture, pretrained on the ImageNet dataset, was used to extract features for use with the shallow model at x5, x10 and x20 magnifications.

### TCGA Data

We initially applied DLVPM to data from the TCGA breast cancer cohort. In order to ensure we only used the highest quality data, we subjected this data to several selection steps prior to its use: Acquisition site can have a strong effect on both omics and imaging data^27^. We used a novel method to remove the effect of acquisition site. However, when a small number of samples are associated with a covariate, it is not possible to disentangle biological effects, and effects driven by this nuisance covariate. Therefore, we only used data from acquisition sites that contributed at least ten samples to the TCGA study. We only used samples with a tumour purity above 60%. This threshold was chosen to minimize contamination from non-cancerous cells, thereby reducing background noise and increasing the precision of genetic and epigenetic profiling. By focusing on samples with higher tumour purity, we aimed to obtain clearer insights into tumour-specific molecular pathways and genetic alterations. We only used female participants.

TCGA data was obtained from: https://portal.gdc.cancer.gov/. A total of 775 patient samples had clinical and SNV data available for the DLVPM-Twins analysis, using the selection criteria specified above. 758 patient samples had a full set of SNV, methylation, miRNA-Seq, RNA-Seq and histological data available for the full path modelling analysis. Training and testing data sets were created in a random 80% and 20% split for the DLVPM Twins and full path modelling analyses. This split was then used consistently through the rest of the investigation.

Histology data was analysed using DLVPM at x5,x10 and x20 magnifications. The first step in that process was to identify which parts of the overall image contained histological tissue. We did this by calculating Sobel’s image gradient across the whole slide. We then split the tissue into 224×224 pixel sections, subsequently referred to as image tiles. This tile size is the input size required by the EfficientNetB0 architecture^29^ used here. This process was repeated at ×5, ×10 and ×20 magnifications. Tiles where the average Sobel’s image gradient was over 15 for over 50 percent of the image were considered to contain enough tissue to be used to train the DLVPM model.

We downloaded Single Nucleotide Variant (SNV), Methylation, miRNA-Seq and RNA-Seq data from the TCGA database. This data is very high dimensional. First, we reduced the data dimensionality for each modality. In the case of the Methylation and the RNA-Seq data, we did this by finding the genes with the top 10 percent highest variance. miRNA-Seq data has much lower dimensionality, so we used the top 50 percent here. Omics data such as RNA-Seq are often heavily skewed. We subjected all omics data to a rank based inverse gaussian transform to remove this data skew.

### Mediation Effects

The effect of DLVs constructed from Methylation, miRNA-Seq and SNV data should all act indirectly on histology, with RNA-Seq acting as a mediator. We tested for mediation effects using the ‘statsmodels’ package. Our mediation model designated the DLVs derived from Methylation, miRNA-Seq, and SNV data as independent variables, RNA-Seq data as the mediator, and histological outcomes as the dependent variable. To assess the significance of the mediation effect, ‘statsmodels’ employs a bootstrapping approach; By using bootstrapping, ‘statsmodels’ does not rely on the assumption of normality for the indirect effect, making it a robust method for mediation analysis. The results of the bootstrapped mediation analysis provided an estimate of the size and significance of the indirect effects of Methylation, miRNA-Seq, and SNV data on histology through RNA-Seq data.

### Individually Significant Effects

DLVPM is a method for identifying global associations between different data types. We carried out additional analyses to localise effects to individual genetic loci. We ran these analyses to determine both the overall significance of individual genetic loci within the path modelling analysis, and the significance of their association with histological data. When calculating the overall significance of individual genetic loci within the path modelling analysis, we used the following procedure: For each of the multi-omic data types, we calculated the mean association between each ‘omics loci in that data-type, and the DLVs that are connected to that data-type via the DLVPM path model, in the testing set.

We then used permutation testing to ascribe significance to the mean of these associations for each multi-omic data type. Using permutation testing, it is possible to correct for multiple comparisons by using the maximal statistic across all loci, in this case the largest mean correlation coefficient, as the statistic of interest in the permutation distribution^82^. This procedure controls the family-wise error in the strong sense.

We used the same procedure when determining the significance of associations between the histology data, and individual genetic loci. The only difference in this analysis was that we only calculated associations between individual genetic loci, and the histology DLVs, rather than taking the mean association as our statistic of interest.

### Gene Set Enrichment Analyses

We carried out a gene set enrichment analysis (GSEA) on the ranking of correlations between gene expression scores, quantified by RNA-Seq, and DLVs from connected datasets (see Methods). Results from these analyses are shown in Extended Data Figure 7. The GSEA analysis was carried out using the fgsea package in R. This analysis used the gene set ‘Hallmarks’, downloaded from: https://www.gsea-msigdb.org/gsea/msigdb/human/collections.jsp. This analysis was conducted using default exclusion criteria, whereby pathways with fewer than 15, or more than 500, genes were omitted from the analysis. Significance was determined using an adjusted p-value threshold of less than 0.1. The normalized enrichment score (NES) was utilized to evaluate effect sizes.

### Histological Visualisation

Our DLVPM model undergoes training using tissue tiles and, upon completion, we deploy it to analyse each tile individually. This enhances our understanding of the tumour segments that exhibit the most pronounced effects for specific Deep Latent Variables (DLVs). This allows us to pinpoint and assess the tumour sub-sections that have the greatest influence on each DLV.

### Single Cell Analysis

Single cell data was obtained from: https://www.ncbi.nlm.nih.gov/geo/query/acc.cgi?acc=GSE176078. We applied the RNA-Seq component of the full DLVPM model, trained on the TCGA dataset to data from the single cell breast cancer encyclopedia^49^. The single cell breast cancer encyclopaedia is a collection of 100,064 single cells with transcriptomic data, taken from 26 primary tumours including 11 ER+, 5 HER2+ and 10 TNBC, representing the three major clinical subtypes. This data was pre-processed in the same manner as the TCGA RNA-Seq data.

### Cell-Line Data

Cell-Line data was obtained from: https://depmap.org/portal/. Of the available breast cancer cell line data, 61 RNA-Seq samples were available, 67 SNV samples, 50 miRNA-Seq samples and 45 CRISPR-Cas9 samples. All of this ‘omics data was used in the analyses presented here. Pairwise associations between ‘omics data-types, and CRISPR-Cas9 data, utilised all intersecting samples.

Omics data from the depmap project was pre-processed for use in the same manner as data from the TCGA dataset. Although methylation data was collected as part of this project, this data is of a different type to that collected as part of the TCGA project. This data was therefore not used as part of the current investigation. In some cases, data was not available for particular genes/loci. These genes/loci were replaced by columns of zeros. The DLVPM model was robust to these changes as it was trained with a dropout layer, simulating this effect.

As noted earlier, we used a confound layer to remove the effects of nuisance covariates when training the DLVPM model on TCGA data. When we applied the model to CCLE data, this layer was removed from the model as these covariates are not relevant for CCLE data. We also used batch level statistics to ensure DLVs were orthogonal in this new dataset.

### Spatial Transcriptomics

Spatial transcriptomic data was obtained from: https://www.10xgenomics.com/. At the time of the analysis, four breast cancer samples were available using the Xenium platform. DLVPM was initially applied to TCGA data to parse inter-tumoral heterogeneity. Because histology data is trained on sections of tissue called tiles, it is possible to deconvolve tile-wide effects back into image space. This allows us to visualize histologic heterogeneity across individual tumours. Recently, a range of spatial transcriptomic methods have been developed with the aim of quantifying heterogeneity in gene expression across individual tumours.

The Xenium platform, from 10X genomics, is an in-situ hybridisation based spatial transcriptomic method^67^. This platform provides sub-cellular transcript resolution for genes known to be important in Breast Cancer. We sought to identify relations between DLVPM models, and genes found to be essential to the functioning of cells scoring highly on these models. The DLVPM histological model has a tile-wise resolution of 224x224 pixels. We extracted tile-wise histological DLVs, and calculated the association between these DLVs, and the total number of transcripts of genes of interest in the matching tile, normalized by the total number of transcripts.

We assessed the significance of associations between DLV 1, and the genes: CCND1, FOXA1, GATA3 and ESR1. As there is a high degree of spatial autocorrelation in this data, uncritical application of Pearson’s coefficient will lead to inflated significance levels, and type-1 errors. For this reason, we used a method to assess statistical significance that fully accounts for spatial autocorrelation ^83^.

## Data Availability

All data used in this investigation is publicly available. TCGA data can be found at: https://portal.gdc.cancer.gov. Data from the single cell breast cancer encyclopaedia can be downloaded from: https://www.ncbi.nlm.nih.gov/geo/query/acc.cgi?acc=GSE176078. Cancer dependency map data can be found at: https://depmap.org/portal/. Spatial transcriptomic breast cancer data derived from the Xenium platform can be downloaded from: https://www.10xgenomics.com/.

## Code Availability

Code implementing the Deep Latent Variable Path Modelling (DLVPM) method can be found at: https://github.com/alexjamesing/Deep_LVPM. All other scripts used for plotting and analysis are available by request from the first author.

## Author Contributions

AI formulated the DLVPM method, wrote the code, carried out all analyses and produced all the figures except the GSEA analysis, which was run by AA. JOK provided scientific direction and supervision. AI, JOK, AA and MC interpreted the data. AI wrote the first draft of the paper, with subsequent contributions from JOK, AA and MC.

---

**Extended Data Figure 1:**
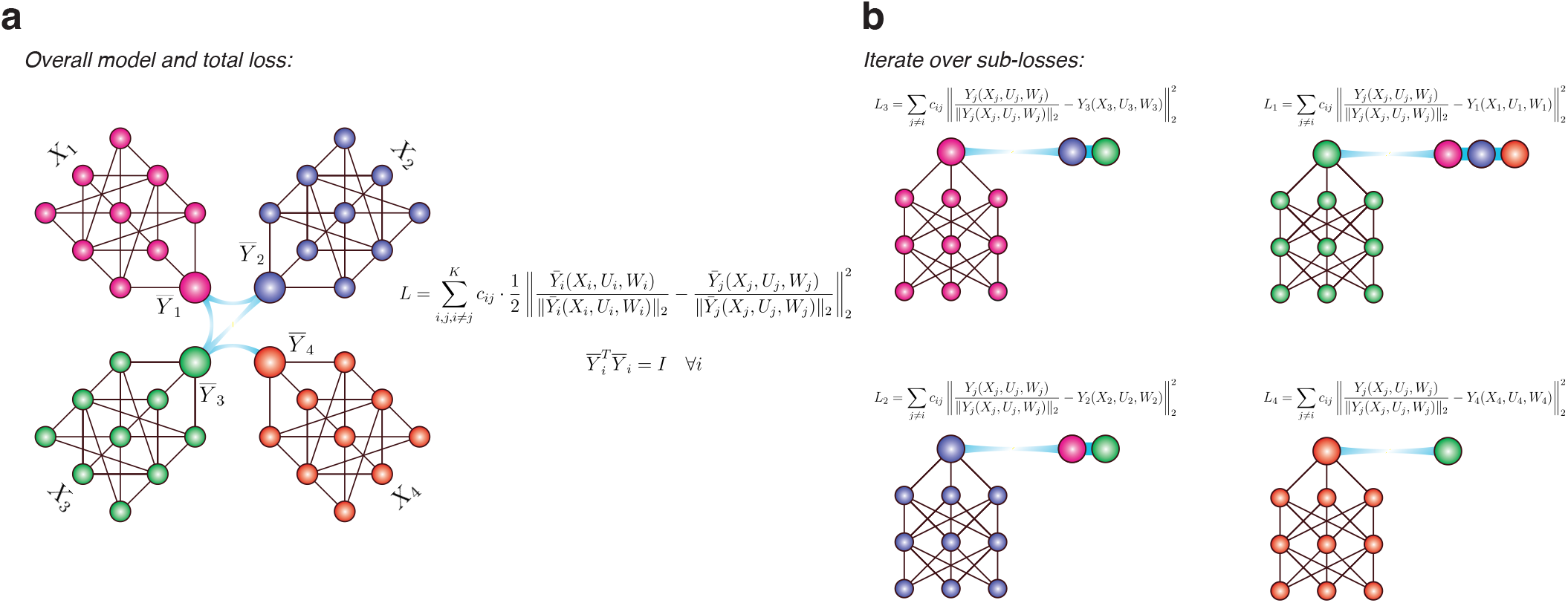
Illustrates the DLVPM model, and the manner in which it is trained. a: Shows the overall model. The aim of the method is to maximize the sum of correlations between deep latent variables (DLVs) connected by the path model. This optimization can be achieved by minimising a sum of least squares losses between the output of each measurement model, and the measurement models they are connected to via the path model. b: The overall loss can be minimized by iteratively minimizing least squared losses between the output of each measurement model, with outputs from connected measurement models.

**Extended Data Figure 2a:**
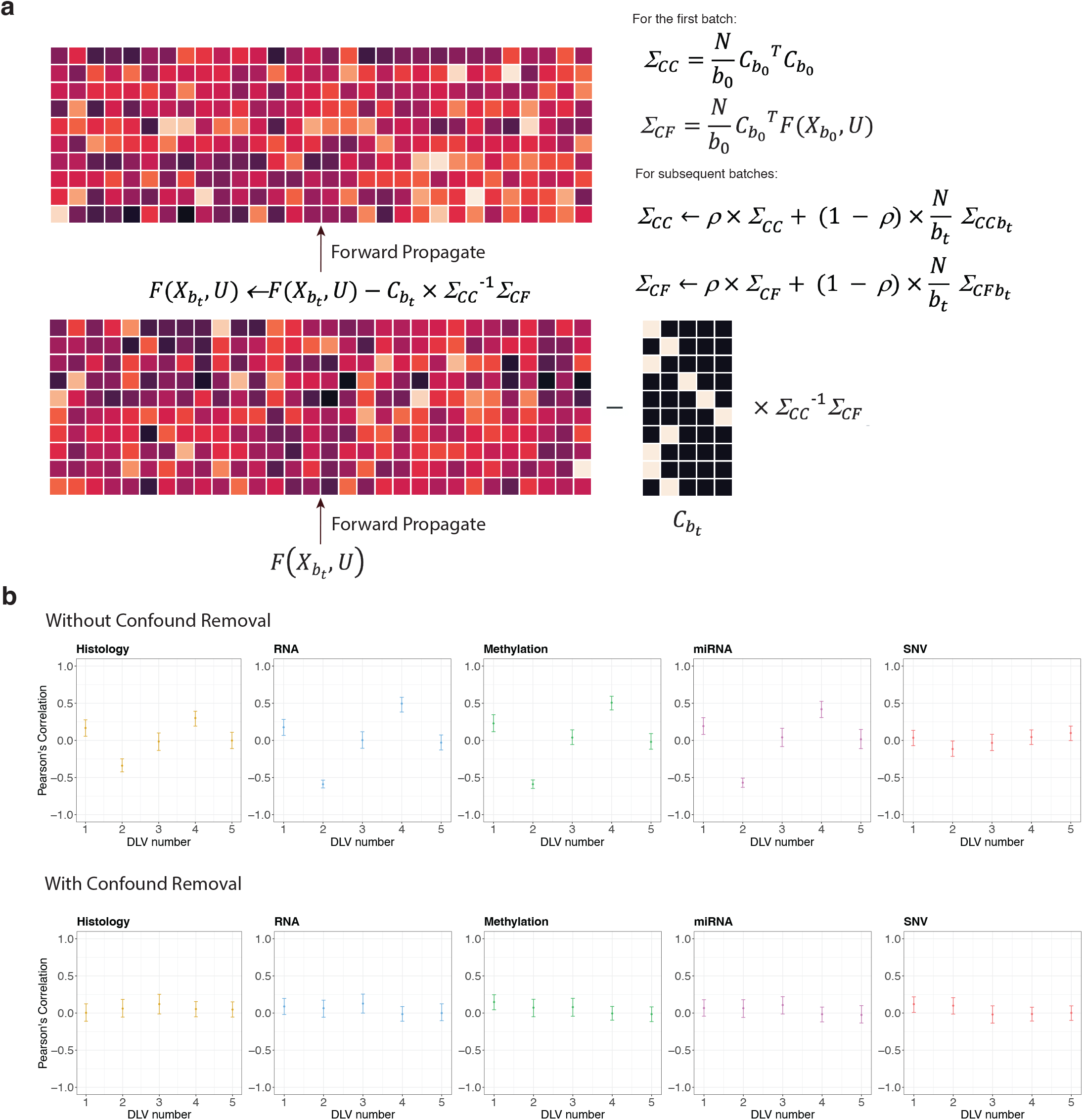
Confound Removal in Deep Latent Variable Projection Models (DLVPM). This panel illustrates the batch-wise confound removal process, where confounding variables are accounted for and orthogonalized in the penultimate layer of the model to ensure that the final output is independent of these variables. The first step involves calculating and subtracting the confound influence from the batch signal, followed by the propagation of the corrected signal through the network. Mathematical definitions of the variables shown here are given in the Methods. b: displays the effectiveness of the confound removal method, contrasting the association between the deep latent variables (DLVs) and a confounding factor before and after the application of the confound removal process across different data types.

**Extended Data Figure 3:**
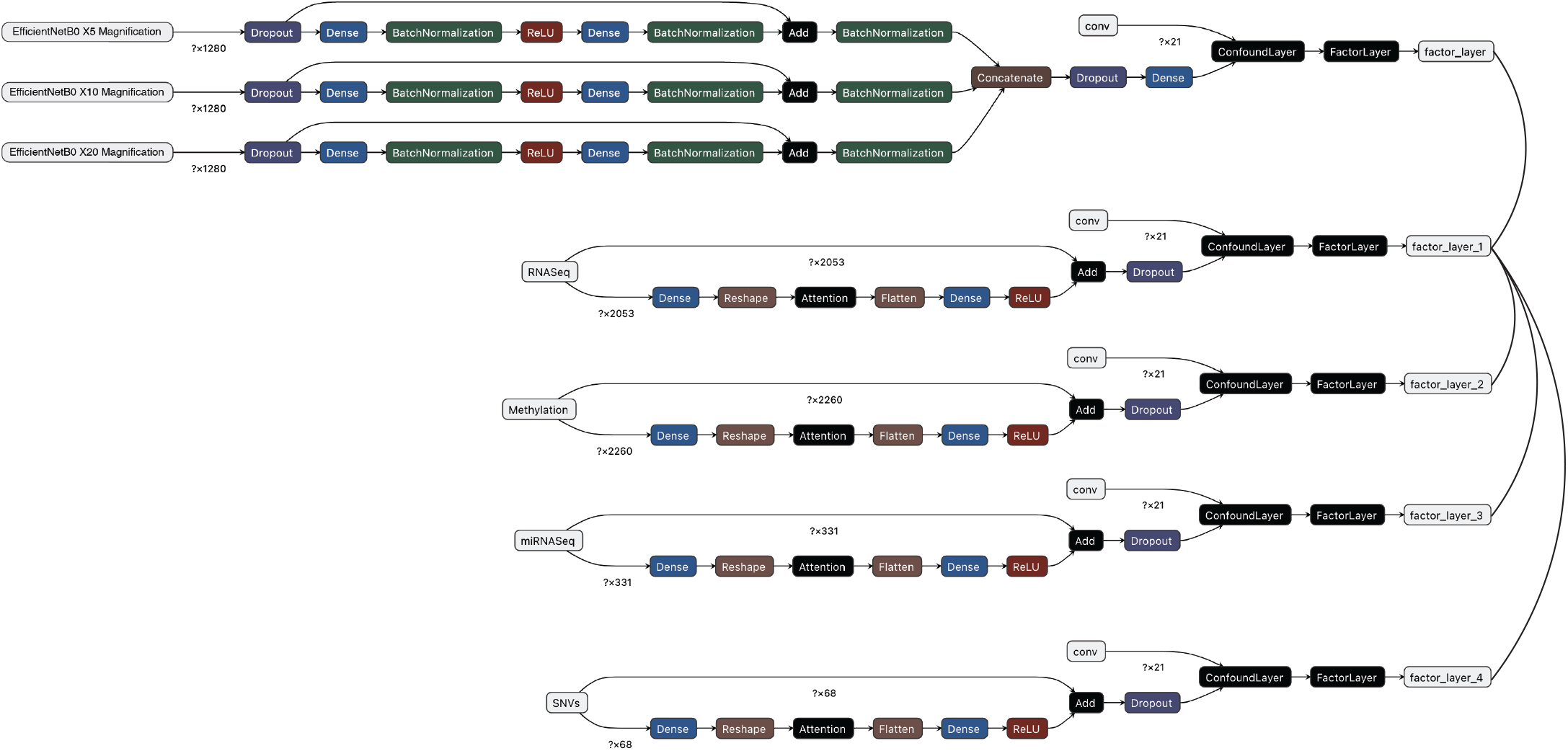
Illustrates the neural network models used as measurement models for each data type in the present investigation. The figure shows the neural network used to process histology data. There are three inputs to the network. These inputs take imaging data that has been tiled, passed through a pre-trained EfficientnetB0 architecture, and then mean averaged over the network output for each tile. The network takes these inputs, and passes them through a feed forward network to produce an output for each magnification, which are then concatenated before DLV estimation. Skip connections are used so that information from lower layers can pass freely through the network. Each of the omics data modalities was processed with a residual network that combines an attentional mechanism with the raw input via a skip connection.

**Extended Data Figure 4a:**
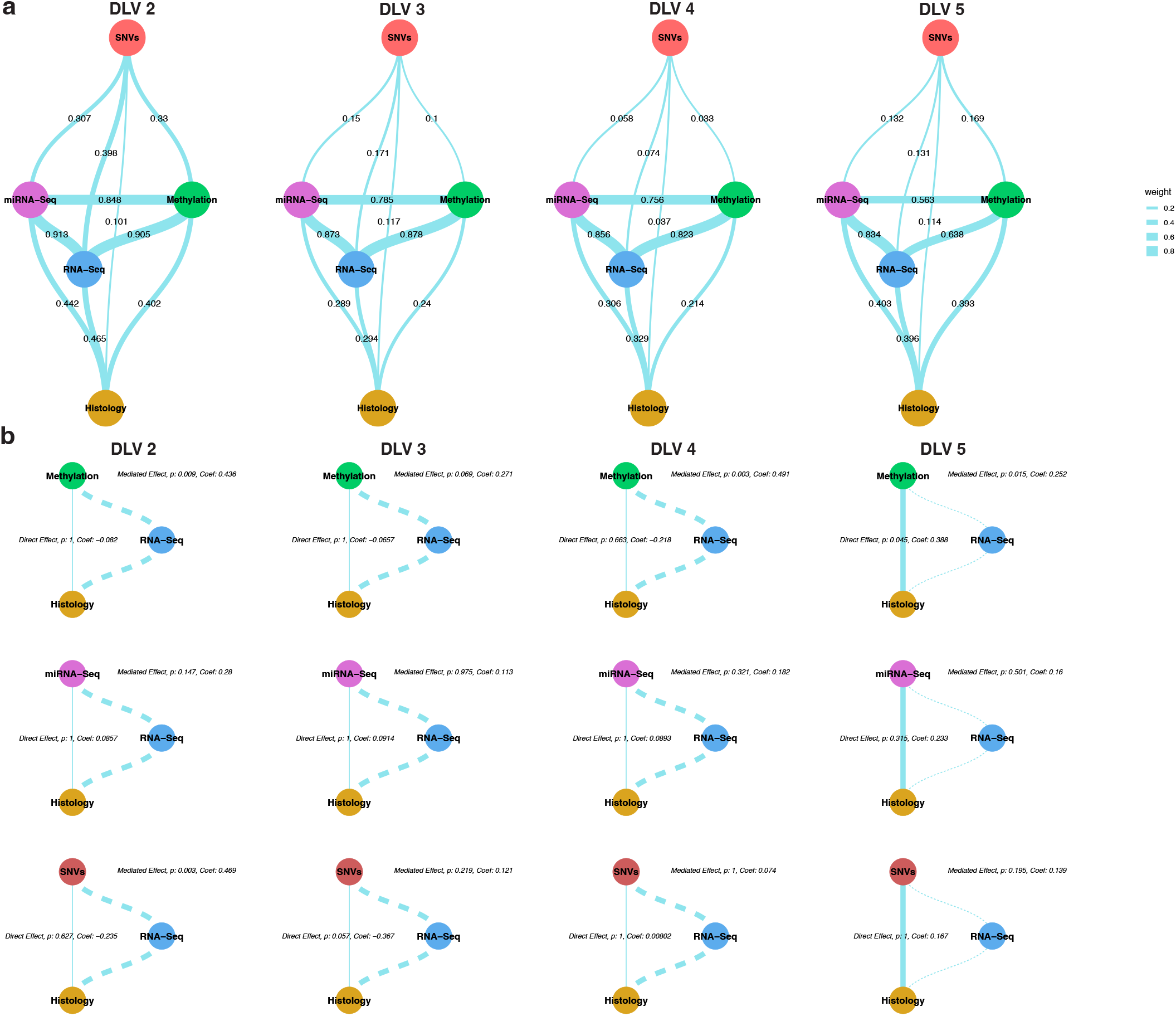
This figure panel shows DLVPM path models for all DLVs after the first, trained and tested on TCGA data. Each of these network graphs represents an orthogonal associative mode linking different data, established using the DLVPM procedure. b: These network graphs illustrate different multi-omic mediation analyses using DLVs estimated using DLVPM. In each case, DLVs from the histology data take the place of dependent variables. DLVs from the RNA-Seq data act as mediator variables. The significance of direct and mediation effects of DLVs derived from: SNVs, miRNA-Seq and Methylation data were all tested in mediation models. Significance of direct and indirect effects was determined using bootstrapping. Straight paths between the independent variable and the dependent variable represent direct effects, where the width of the edge linking them is equal to the beta value for the direct effect in the mediation model. The indirect effect is represented by the path through the RNA-Seq DLV.

**Extended Data Figure 5:**
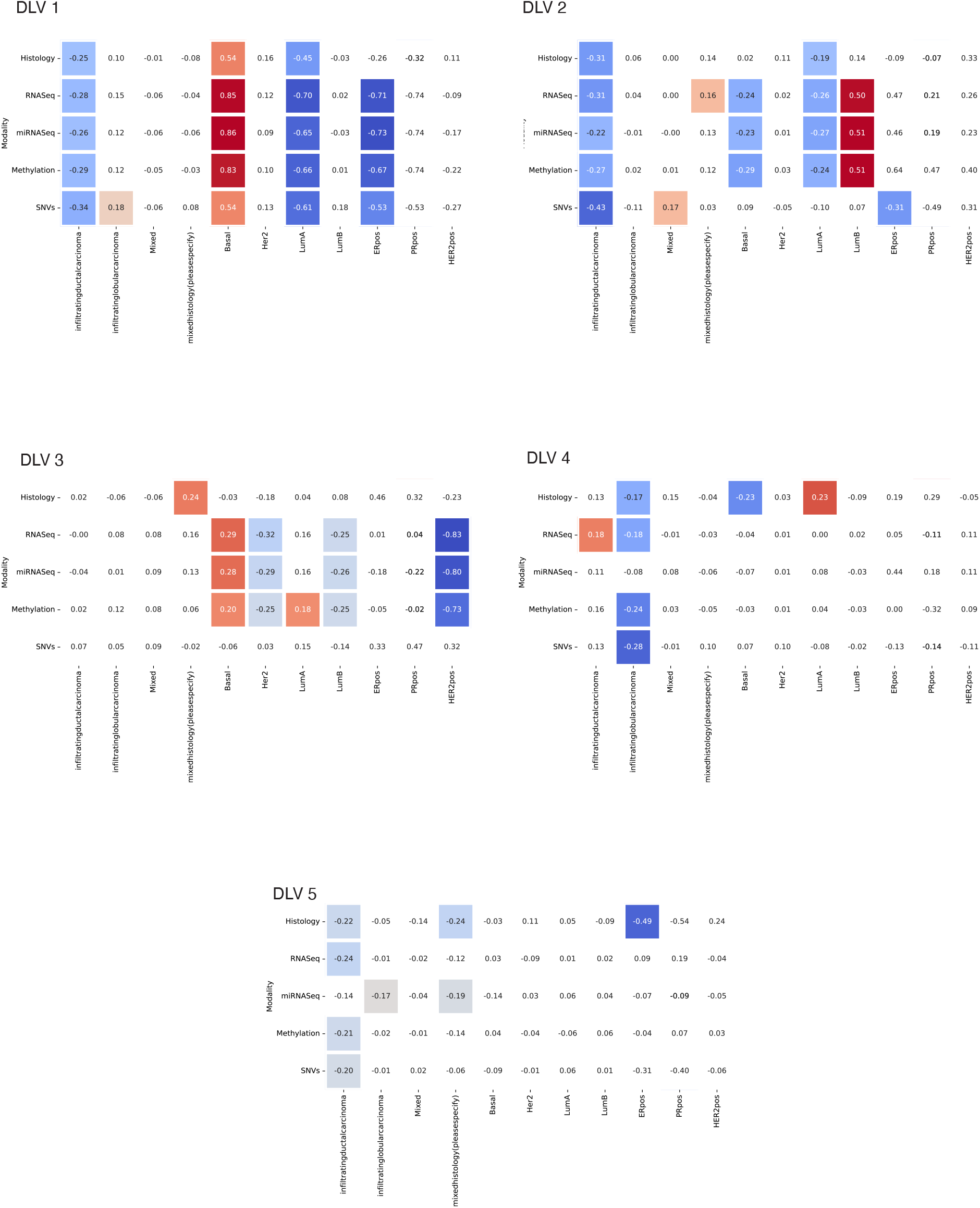
The heat maps shown in this figure illustrate associations between DLVs, estimated using the full DLVPM model, and clinical molecular and histological characteristics. Quantitative values in each heatmap are the Pearson’s Correlation Coefficient between different clinical types, and DLVs. Heatmap squares that are coloured are significant at the p < 0.05 level. Squares that are not coloured are non-significant.

**Extended Data Figure 6:**
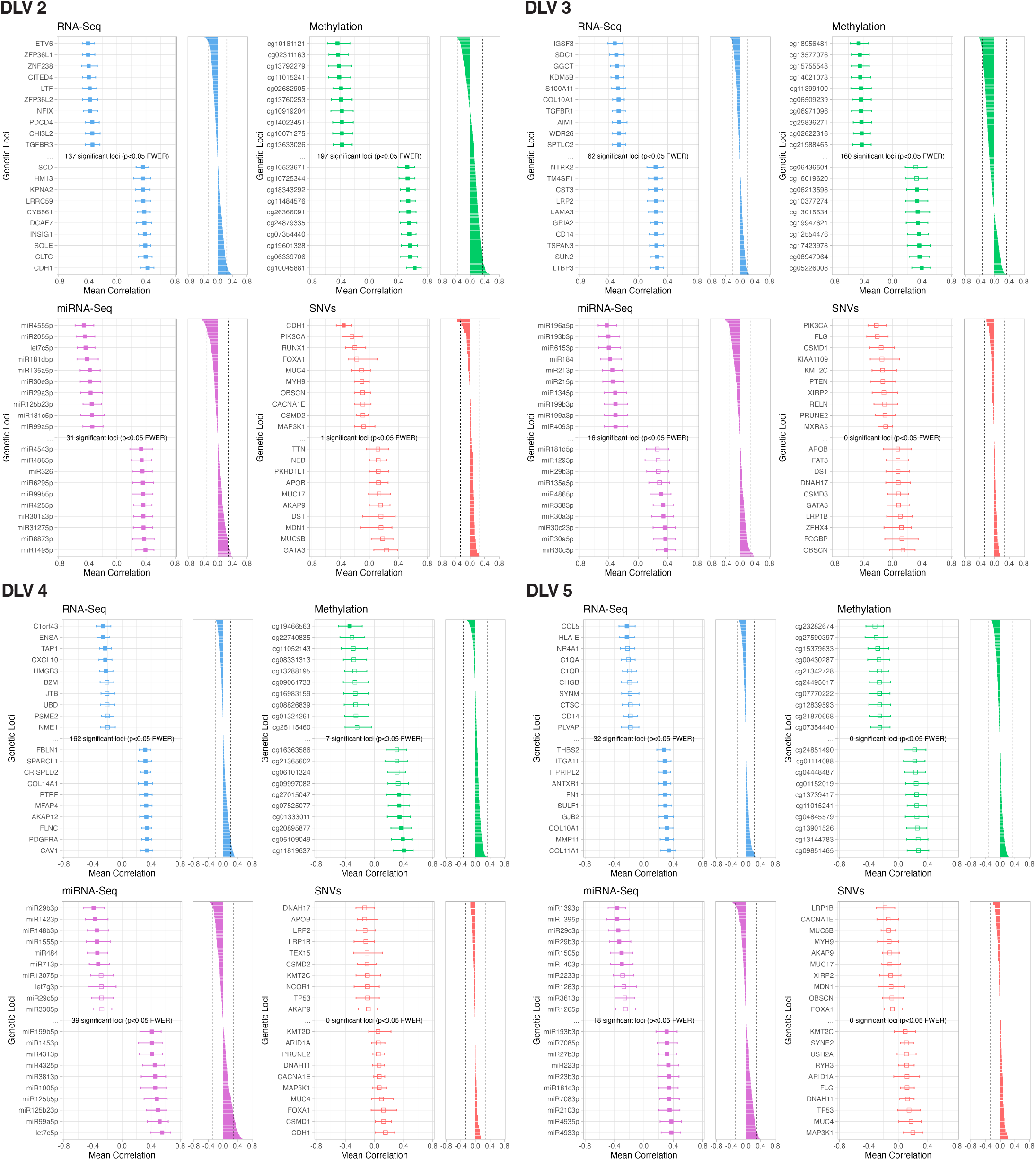
Results of additional analyses to localize effects to particular genetic loci. The plot shows association values between genetic loci, and DLVs connected to the data-view under analysis by the path model. The panel on the left shows the ten most positively and negatively associated genetic loci. The error bars represent 95% bootstrapped confidence intervals. The panel on the right shows all genetic loci under analysis along with the significance threshold cut-off

**Extended Data Figure 7:**
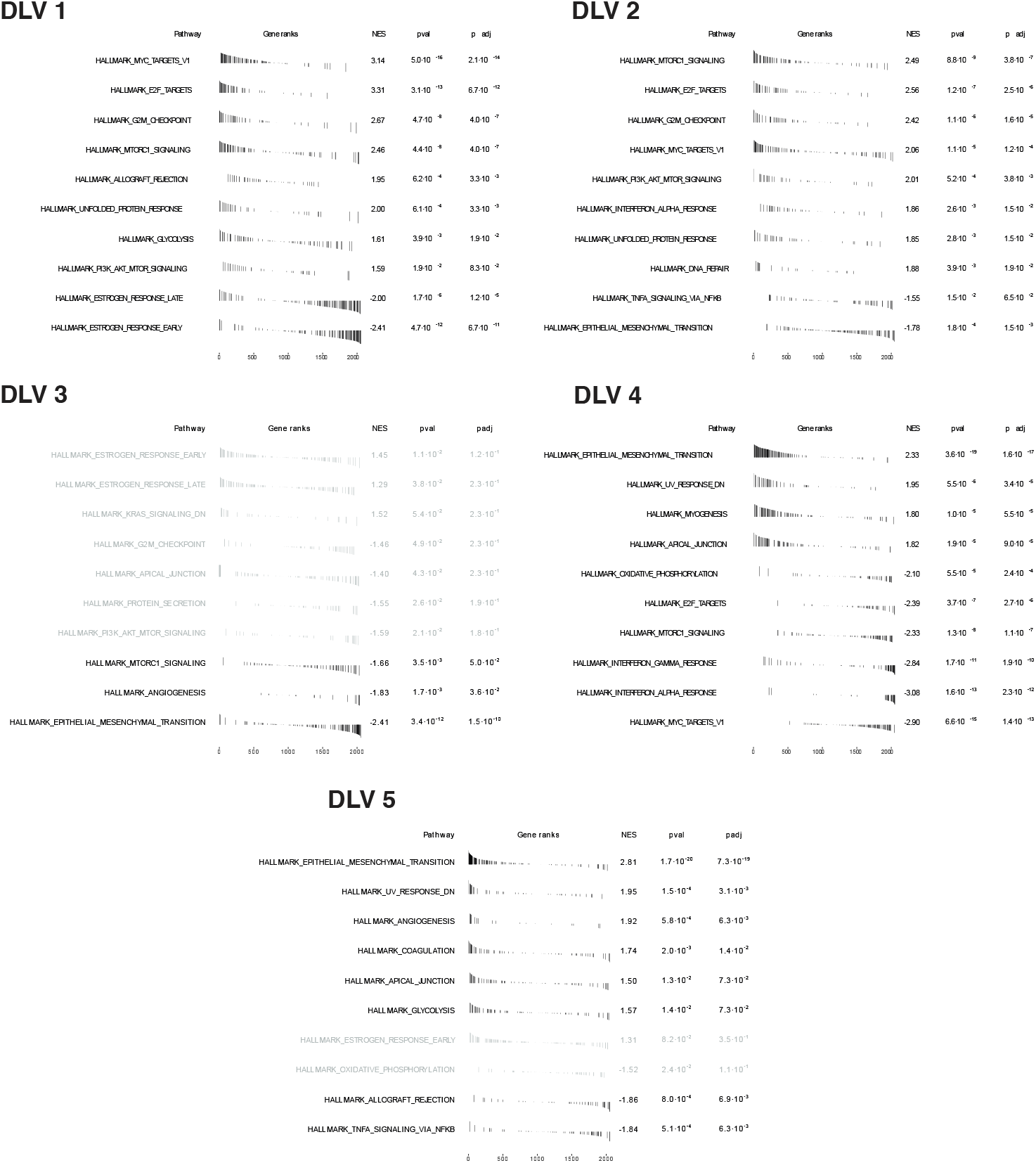
This figure shows the top 10 most positively enriched, and top 10 most negatively enriched ‘hallmarks of cancer’ ontology terms for each of the DLVs extracted by DLVPM. Terms that are non-significant at padj > 0.1 have been greyed out.

**Extended Data Figure 8:**
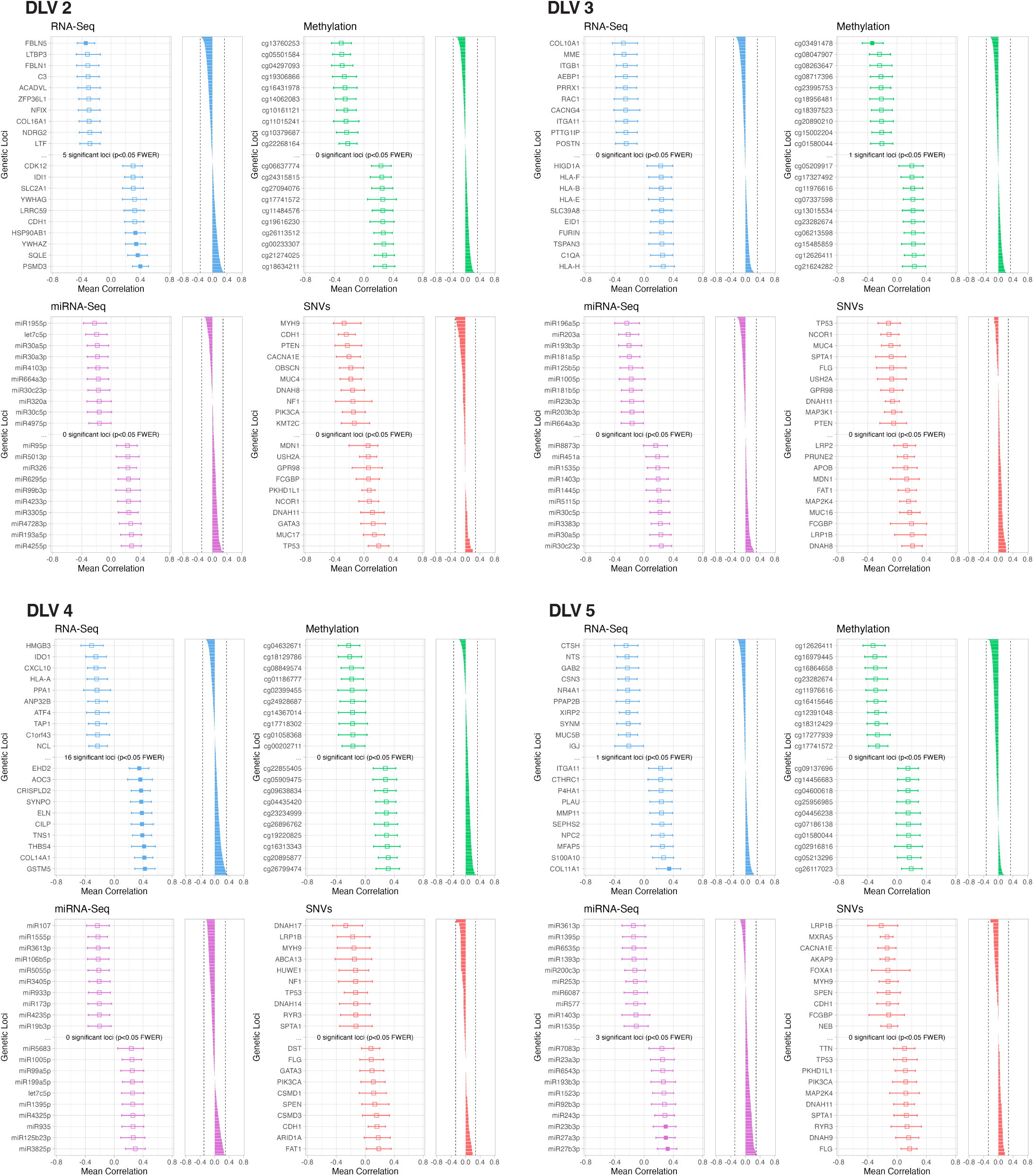
The plot shows association values between genetic loci, and DLVs connected to the data-view under analysis by the path model. The panel on the left shows the ten most positively and negatively associated genetic loci. The error bars represent 95% bootstrapped confidence intervals. The panel on the right shows all genetic loci under analysis, along with the significance threshold cut-off.

**Extended Data Figure 9:**
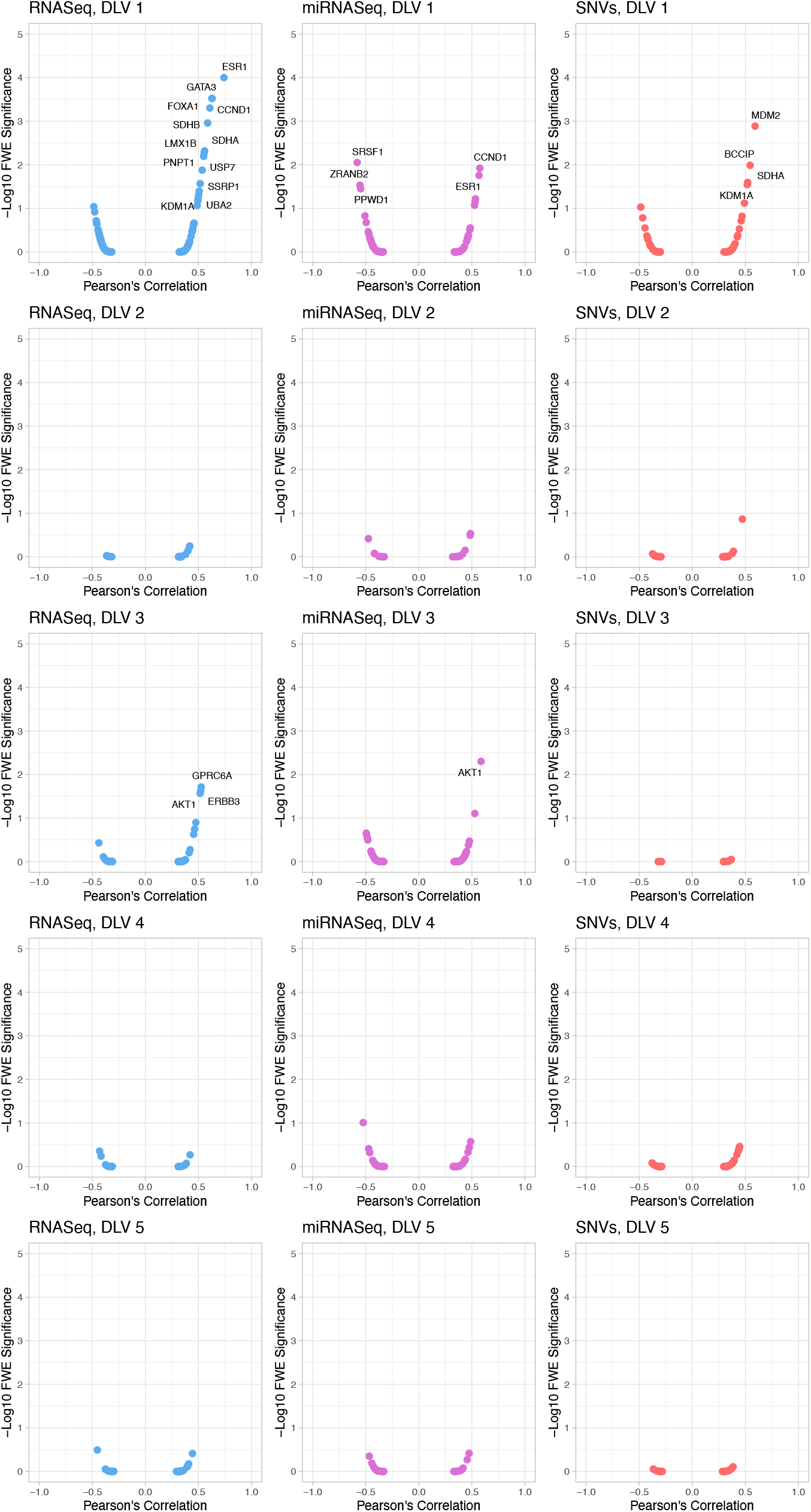
These figures show plots of pairwise association of DLVs against -log10 p-values, plotting the relation between DLVs and gene dependency scores derived from RNAi data.

**Extended Data Figure 10:**
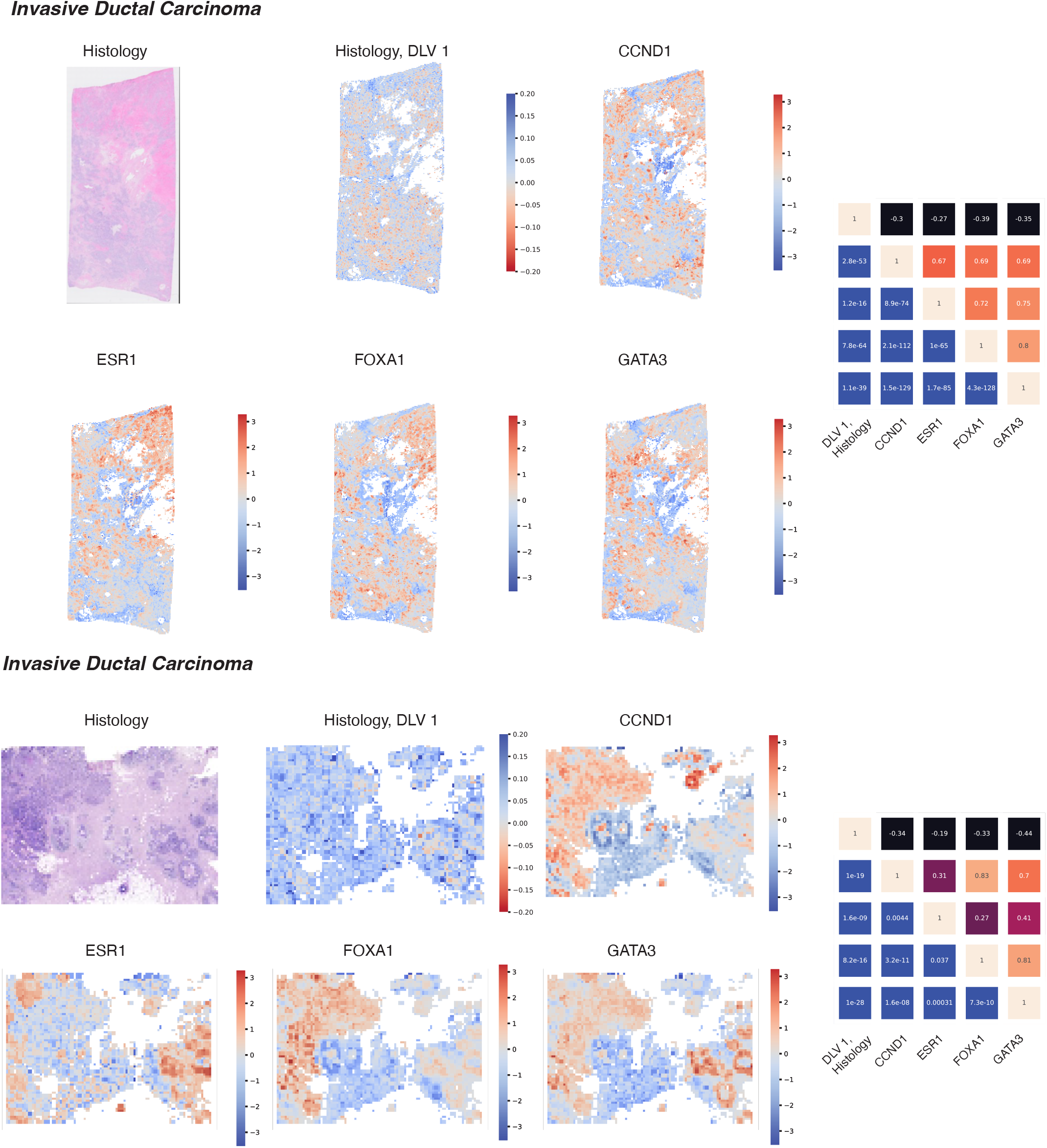
Tile-wise heatmaps generated from the DLVPM model, trained on TCGA data, and applied to histological and associated spatial transcriptomic data. The colormap is flipped for the histology heatmap as this DLV shows a negative association with the genes of interest. We applied this analysis to invasive ductal carcinoma and invasive lobular carcinoma. The association/significance matrices on the right show correlations between genes of interest and the first histology DLV for both tumours. The upper triangular part of each matrix is denoted with Pearson’s Correlation Coefficient between each gene, and the histology data. The lower triangular part of each matrix denotes the significance level between genes and histology data.

## Notes

https://github.com/alexjamesing/Deep_LVPM

### Competing Interest Statement

The authors have declared no competing interest.

