## Supplementary Information for "Integrating Multi-Modal Cancer Data Using Deep Latent Variable Path Modelling"

The supplementary information contains only table captions for Supplementary Table 1 and Supplementary Table 2.

*Supplementary Table 1: Sheets from this table contain mean Pearson’s Correlation values between each genetic loci, and DLVs that modality is connected to via the path model. DLVs are labelled from DLV1 to DLV5. Each of the mean correlation values also has a Family-wise error corrected (FWER) significance level attached.*

*Supplementary Table 2: Sheets from this table contain Pearson’s Correlation values between each genetic loci, and the histological DLVs. DLVs are labelled from DLV1 to DLV5. Each of the mean correlation values also has a Family-wise error corrected (FWER) significance level attached.*
